# Environmental DNA phylogeography: successful reconstruction of phylogeographic patterns of multiple fish species from a cup of water

**DOI:** 10.1101/2022.09.02.506317

**Authors:** Satsuki Tsuji, Naoki Shibata, Ryutei Inui, Ryohei Nakao, Yoshihisa Akamatsu, Katsutoshi Watanabe

**Affiliations:** Graduate School of Science, Kyoto University, Kitashirakawa-Oiwakecho, Sakyo-ku, Kyoto 606–8502, Japan; Graduate School of Science and Technology for Innovation, Yamaguchi University, 2-16-1 Tokiwadai, Ube, Yamaguchi, 755-8611, Japan; Environmental Research and Solutions co., ltd, Hikaridai 2–3–9, Seika-cho Sourakugun, Kyoto 619–0237, Japan; Faculty of Socio-Environmental Studies, Fukuoka Institute of Technology, Wajiro-higashi, Higashi-ku, Fukuoka 811–0295, Japan

**Keywords:** comparative phylogeography, environmental DNA, freshwater fish

## Abstract

Phylogeography is an integrative field of science linking micro- and macro-evolutionary processes, contributing to the inference of vicariance, dispersal, speciation, and other population-level processes. Phylogeographic surveys usually require considerable effort and time to obtain numerous samples from many geographical sites covering the distribution range of target species; this associated high cost limits their application. Recently, environmental DNA (eDNA) analysis has been useful not only for detecting species but also for assessing genetic diversity; hence, there has been growing interest in its application to phylogeography. As the first step of eDNA-based phylogeography, we examined (1) data screening procedures suitable for phylogeography and (2) whether the results obtained from eDNA analysis accurately reflect known phylogeographic patterns. For these purposes, we performed quantitative eDNA metabarcoding using group-specific primer sets in five freshwater fish species belonging to two taxonomic groups from a total of 94 water samples collected from western Japan. As a result, three-step data screening based on the DNA copy number of each haplotype detected successfully eliminated suspected false positive haplotypes. Furthermore, eDNA analysis could almost perfectly reconstruct the phylogenetic and phylogeographic patterns obtained for all target species with the conventional method. Despite existing limitations and future challenges, eDNA-based phylogeography can significantly reduce survey time and effort and is applicable for simultaneous analysis of multiple species in single water samples. eDNA-based phylogeography has the potential to revolutionise phylogeography.

## Introduction

Since its emergence in 1980s, the field of phylogeography has rapidly developed as an integrative field of science linking micro- and macro-evolutionary processes (Avise, 2000; Avise et al., 1987; Bermingham and Moritz, 1998; Soltis et al., 2006). Phylogeography explores the historical and ecological processes and principles shaping the geographical distribution patterns of genealogical lineages within a species or closely related species by inferring vicariance, dispersal, speciation and other population-level processes (Avise et al., 1987; Sersics et al., 2011). Furthermore, comparative phylogeography, which focuses on comparing phylogeographical structures among co-distributed species, allows us to detect biodiversity patterns at regional levels (Bermingham and Moritz, 1998). Moreover, it contributes to the understanding of the broad impacts of geological events or environmental conditions on the distribution range and demographic history of multiple species, predicting the responses of biological populations to climate change (Moritz, 2002; Scoble and Lowe, 2010; Watanabe et al., 2006).

Despite the high importance of phylogeography, much effort and time are usually required to uncover species’ phylogeographic patterns. To investigate a phylogeographic structure, a few dozen of individuals (usually 10–40 individuals) from each local population of the target species should be captured to obtain specimens or tissue samples. This requirement is one of the major barriers to phylogeographic studies. For example, when targeting mobile aquatic species, sampling sometimes takes years or decades to complete (Corush et al., 2022; Miyake et al., 2021; Nakagawa et al., 2016; Ruzzante et al., 2008). Additionally, capture surveys may be limited by safety considerations, laws and/or the potential invasiveness of tissue collection, which may raise various ethical considerations (Dugal et al., 2022; Tsuji et al., 2020b). Therefore, the development of cost-effective and non-invasive approaches to overcome these limitations may facilitate phylogeography studies.

Recently, environmental DNA (eDNA) analysis has been demonstrated as highly useful not only for detecting species diversity but also for assessing genetic diversity; leading to growing interest in its application to phylogeography (Parsons et al., 2018; Shum and Palumbi, 2021; Sigsgaard et al., 2020; Tsuji et al., 2020a; Turon et al., 2020). The application of eDNA analysis would revolutionise the way to assess the phylogeographic structure for at least three reasons. First, eDNA is DNA material released from organisms into the environment (e.g. soil, water and air); as such, eDNA surveys only require the collection of environmental samples instead of capturing the target species (Taberlet et al., 2012; Thomsen and Willerslev, 2015). This minimises the effort and time required for field surveys, virtually eliminating the damage to target species and their habitats. Second, a single eDNA sample contains information from many individuals from local populations of various species at the sampling site; therefore, the same eDNA sample can be used to assess genetic diversity in multiple species by using a group-primer set (i.e. primers for simultaneous DNA amplification from a related species group) or by changing the species-specific primer set (Shum and Palumbi, 2021; Tsuji et al., 2020c; Turon et al., 2020; Weitemier et al., 2021). Third, sampling universality and the high preservation of eDNA samples under freezing conditions allow reusing previously collected samples for other studies (Andres et al., 2021; Yamamoto et al., 2017; Zizka et al., 2022). This would save sampling time and effort for new studies.

To applyeDNA analysis to phylogeography, we first need to examine whether the geographical distribution patterns of genetic lineages estimated by eDNA analysis accurately reflect the results from the conventional method using tissue DNA and Sanger sequencing. Previous studies have so far focused primarily on whether eDNA-obtained haplotypes could be assigned to known haplotypic variations (e.g. Baker et al., 2018; Holman et al., 2022; Sigsgaard et al., 2016; Stat et al., 2017; Yoshitake et al., 2019). Some studies have attempted to detect region-specific haplotypes, but the number of study sites was limited or there was no available phylogeographic information obtained with the conventional method (Nguyen et al., 2021; Parsons et al., 2018; Turon et al., 2020; Weitemier et al., 2021). In addition, for haplotype detection based on eDNA analysis, it is essential to properly remove erroneous sequences derived from high-throughput sequencing data (Tsuji et al., 2020b; Turon et al., 2020). Although several denoising strategies have been proposed, no previous studies have investigated data screening procedures to eliminate false positive sequences and recover more accurate phylogeographic information by eDNA analysis (Parsons et al., 2018; Sigsgaard et al., 2016; Tsuji et al., 2020a).

This study aimed to examine the performance of eDNA analysis as a tool for phylogeography by comparing its results with those of the conventional method. Here, as a model case, we targeted five freshwater fish species (i.e. two odontobutid gobies, *Odontobutis obscura* and *Odontobutis hikimius* and three cyprinids, *Nipponocypris temminckii, Nipponocypris sieboldii* and *Zacco platypus*, all distributed in western Japan) for which detailed phylogeographic information is available or obtained presently using Sanger sequencing of tissue DNA. We specifically examined the following two points: (1) data screening procedures to eliminate false positive sequences and improve the detection reliability of genetic lineages and (2) after appropriate data screening, how accurately the geographical distribution patterns of genetic lineages estimated by eDNA analysis reflect the results of Sanger sequencing-based studies. Accordingly, we discuss the usefulness, potential and current limitation of eDNA-based phylogeography.

## 2 Materials and methods

### 2.1 Target species

We targeted the following five species from two teleost families: *Odontobutis obscura* and *O. hikimius* (Odontobutidae) and *Nipponocypris temminckii, N. sieboldii* and *Zacco platypus* (Cyprinidae) (Fig. 1). Among them, *O. obscura, N. temminckii* and *Z. platypus* are commonly found in the middle of western Japan rivers (Hosoya, 2019). *Nipponocypris sieboldii* is also widely distributed in western Japan, but its habitat has been rapidly declining in recent years; thus, the species is included on the Red List in several administrative areas (Kyoto, Nara, Osaka, Kagawa, Yamaguchi prefectures; Hosoya, 2019). *Odontobutis hikimius* is found only in some rivers flowing into the Sea of Japan in the Shimane and Yamaguchi prefectures and was once recognised as the ‘*O. obscura* Hikimi group’ (Iwata and Sakai, 2002). The phylogeographic patterns for each species have been studied in detail based on conventional, tissue-based Sanger sequencing; *O. obscura* and *O. hikimius* (Mukai and Nishida, 2003; Sakai et al., 1998, using allozyme polymorphisms); *N. temminckii* (Taniguchi et al., 2021); *Z. platypus* (Kitanishi et al., 2016). For *N. sieboldii*, its phylogeographic pattern in central Japan was newly estimated based on the conventional method in this study.

**Fig. 1.**
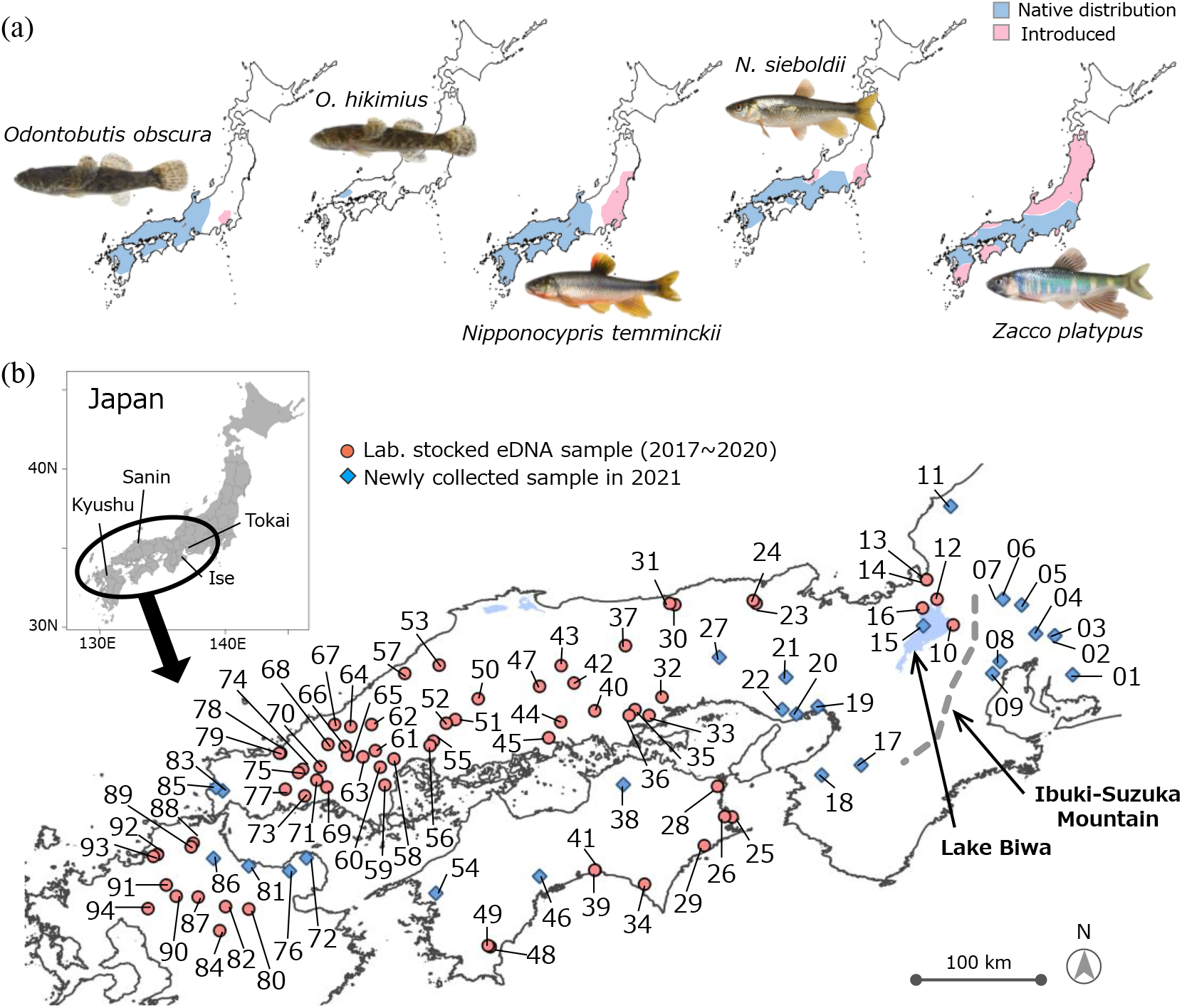
(a) Approximately distribution ranges of each target species in Japan and (b) eDNA sampling locations. Pink circles and blue diamonds indicates samples collected between 2017-2020 and stored in the laboratory and newly collected sample in 2021, respectively. Photo copyright: *O. obscura* and *O. hikimius* for Mr. S. Kunumatsu; *N. temminckii, N. sieboldii* and *Z. platypus* for ffish.asia (https://ffish.asia, 2022.06.01 downloaded).

### 2.2 Group-specific primers and standard DNA development

We developed group primers for species group 1 (*O. obscura* and *O. hikimius*; ‘Odon primer’) and group 2 (*Nipponocypris temminckii, N. sieboldii* and *Z. platypus*; ‘NipZac primer’). First, mitochondrial DNA 12S rRNA (for group 1) and D-loop (for group 2) sequences of target species and closely related species were downloaded from the National Center for Biotechnology Information (NCBI) database (https://www.ncbi.nlm.nih.gov/). For group 1, the same DNA region as in the previous study was selected (Mukai and Nishida, 2003). For group 2, the D-loop region which has a higher mutation rate compared with the other mtDNA regions was selected because it was difficult to design primers meeting the design requirements for cytochrome b (cyt*b*) and NADH dehydrogenase subunit 2 (ND2), targeted in the previous study (Kitanishi et al., 2016; Taniguchi et al., 2021). The downloaded sequences of each region were aligned using MAFFT version 7 (https://mafft.cbrc.jp/alignment/software/).

Based on the aligned sequences, each group-specific primer set was manually designed to contain >1 specific bases within the five bases at the 3′ end of both the forward and reverse primers (Fig. S1). However, for designing the NipZac primers, we could not find any specific SNPs that exclude DNA amplification of *Opsariichthys uncirostris*, which is closely related to group 1 species and originally distributed only in the Lake Biwa–Yodo River system and Mikata Lake, but introduced to other areas in Japan. In fact, the NipZac primers amplified DNA from *O. uncirostris*, but their sequence data did not affect the result as it can be excluded from the data by bioinformatic analysis based on the sequence (Fig. S2a). For primer design, we considered unconventional base pairing in the T/G bond to enhance primer annealing to the template without requiring degenerate bases (Miya et al., 2015). The specificities of the designed group-specific primers were tested *in silico* (Primer-BLAST with default settings; https://www.ncbi.nlm.nih.gov/tools/primer-blast/). Additionally, we performed PCR using designed group-specific primer sets and tissue DNA extracted from the target species of each group and the non-target species considered when designing each primer (three individuals per species, 25 pg/μL per PCR reaction; Fig. S1). PCR conditions were set the same as for the first-round PCR (see section 2.5) for both primer sets. DNA amplification was confirmed by 2% agarose gel electrophoresis.

To quantify the eDNA from each target taxon using the qMiSeq approach (Ushio et al., 2018), we developed standard DNA sets for the respective group-specific primer sets. A consensus DNA sequence of amplification ranges was obtained per group using the sequences of the target taxa used during primer design and Gene Doc software 2.7 (https://genedoc.software.informer.com/2.7/). For each group, two consensus regions (30 base pairs for each) without mutations within taxa were selected and randomly base-changed. The GC% within the selected consensus regions was not changed. Three standard DNAs with different two random regions were designed for each group, respectively (Table S1). The designed standard DNAs were cloned into pEX-A2J2 Vector (Eurofins, Tokyo, Japan). After cloning, the plasmids were cleaved using restriction enzymes and purified using electrophoresis and NucleoSpin Gel & PCR Clean-up kit (Takara bio, Shiga, Japan). The copy number of each standard DNA was calculated based on the concentration quantified with the Qubit 3 Fluorometer (Thermo Fisher Scientific, MA, USA). Finally, standard DNA mixes containing each standard DNA at the following concentrations were prepared for each group: Std. 1 (5 copies/μL), Std. 2 (25 copies/μL) and Std. 3 (50 copies/μL).

### 2.3 Study sites and eDNA collection

Water sampling for eDNA collection was conducted on a total of 94 sites between 2017 and 2021. For 67 of these sites, we used eDNA samples collected for other studies conducted between 2017 and 2020 and stored at –20ºC in the Research Center for Environmental DNA at Yamaguchi University. Samples from the remaining 27 sites were newly collected in 2021 for this study. Details of sample information are shown in Fig. 1b and Table S2.

At each sampling site, we collected 1 L of surface water using a bleached bottle or disposable plastic cup and added benzalkonium chloride (1 mL, 10% w/v; Fujifilm Wako Pure Chemical Corporation, Osaka, Japan) to preserve eDNA (Yamanaka et al., 2017). For the 67 sites surveyed between 2017 and 2020, the collected water samples were transported to the laboratory under refrigeration and filtered using a GF/F glass fibre filter (diameter: 47 mm, mesh size: 0.7 μm; GE Healthcare Japan, Tokyo, Japan) within 36 h after sampling. For the remaining 27 sites surveyed in 2021, the water sample was filtered on-site using a fibre filter. After filtration, all filter samples were immediately stored at –20ºC.

### 2.4 DNA extraction from the filter samples

DNA extraction from filter samples was performed according to two different procedures: Tsuji et al.’s (2022a) method for the 67 sites surveyed between 2017 and 2020; and Tsuji et al.’s (2022b) method for the 27 sites surveyed in 2021. The two procedures described in Tsuji et al. (2022a, 2022b) differed slightly in several respects such as the type of column used to infiltrate the filter into the DNA extraction reagent mix, and the total volume and composition ratio of the extraction reagent mix. In both procedures, the DNA was purified using the DNeasy blood tissue kit and finally eluted in 100 μL Buffer AE. The extracted DNA was stored at –20°C (see appendix for details).

### 2.5 Paired-end library preparation and quantitative eDNA metabarcoding

To construct the paired-end libraries for MiSeq (Illumina, San Diego, CA, USA), a two-step tailed PCR approach was employed. A first-round PCR (1st PCR) was carried out for each of the two groups. The sequences of each group-specific primer combined sequencing primers (italic) and six random hexamers (N) are as follows: Odon_12S primer-F (5′-*ACA CTC TTT CCC TAC ACG ACG CTC TTC CGA TCT* NNN NNN TAT ACG AGA GGC TCA AGC TGA T-3′), Odon_12S primer-R (5′-*GTG ACT GGA GTT CAG ACG TGT GCT CTT CCG ATC* TNN NNN NGT TTT ACC AGT TTT GCT TAC TAT GG-3′); NipZac_D-loop primer-F (5′-*ACA CTC TTT CCC TAC ACG ACG CTC TTC CGA TCT* NNN NNN ACT ATC TTC TGA TAG TAA CCT ATA TGG TA-3′), NipZac_D-loop primer-R (5′-*GTG ACT GGA GTT CAG ACG TGT GCT CTT CCG ATC* TNN NNN NTT GTG TCC CTG ATT CTA TCA TGA ATA G-3′). The insert length in the amplification range of each primer set was ca. 370 bp (group 1, 12S) and ca. 270 bp (group 2, D-loop). The 1st PCR was performed in a 12-μL total volume of reaction mixture containing 6.0 μL of 2 × KAPA HiFi HotStart ReadyMix (KAPA Biosystems, MA, USA), 0.72 μL of forward and reverse primer (10 μM), 2.56 μL of sterilised distilled H_2_O, 1.0 μL standard DNA mix and 1.0 μL eDNA template. The 1st PCR was performed using four replicates per eDNA sample. Additionally, in all 1st PCR runs, four replicates of no-template control using ultrapure water instead of the template and standard DNA mix were prepared and treated as PCR negative control. The thermal conditions for the 1st PCR were 5 min at 95°C, 45 cycles of 20 s at 98°C, 20 s at 60°C, 40 s at 72°C and 5 min at 72°C. The four 1st PCR product replicates were pooled and purified using Sera-Mag SpeedBeads Carboxylate-Modified Magnetic Particles (Hydrophobic) (Cytiva, MA, USA) (target amplicon length including 1st-PCR primers; ca. 500 bp for group 1, ca. 404 bp for group 2). The purified 1st PCR products for each group were adjusted to 0.1 ng/μL and mixed in equal quantities (hereafter ‘1st PCR product mix’) to use as second-round PCR template (2nd PCR).

The 2nd PCR was performed in a 12-μL total volume of reaction mixture containing 6.0 μL of 2 × KAPA HiFi HotStart ReadyMix, 2.0 μL of each primer with index (1.8 μM) and 2.0 μL of the purified 1st PCR product mix. The sequences of the 2nd PCR primers with adapter sequences (underline), eight indexes (X; Hamady et al., 2008) and sequencing primers (italic) were as follows: forward primer (5′-<u>AAT GAT ACG GCG ACC ACC GAG ATC TAC A</u>XX XXX XXX *ACA CTC TTT CCC TAC ACG ACG CTC TTC CCA TCT*-3′); reverse primer (5′-<u>CAA GCA GAA GAC GGC ATA CGA GAT</u> XXX XXX XX*G TGA CTG GAG TTC AGA CGT GTG CTC TTC CGA TCT*-3′). The 2nd PCR primers were used in unique combinations for each sample. The thermal conditions for the 2nd PCR were 3 min at 95°C, 12 cycles of 20 s at 98°C, 15 s at 72°C and 5 min at 72°C. The indexed 2nd PCR products were pooled, and each target band (ca. 580 bp for group 1, ca. 484 bp for group 2) was excised using a 2% E-Gel SizeSelect Agarose Gels (Thermo Fisher Scientific). The sequence library was adjusted to 10 pM (assuming 1 bp DNA has a molecular weight of 660 g/mol) and sequenced on the MiSeq platform at the Environmental Research and Solutions co., ltd (v2 Reagent Kit for 2 × 250 bp PE cartridge, 5% PhiX spike-in). All raw sequences were deposited in the DDBJ Sequence Read Archive (accession number: DRA014749).

### 2.6 Bioinformatic analysis and DNA copy number estimation

The denoising the FASTQ data, the conversion of the number of sequence reads to the DNA copy number and species assignments were performed separately for each group as follows: FASTQ files containing raw reads were denoised using the Divisive Amplicon Denoising Algorithm 2 (DADA2) package version 1.22 (Callahan et al., 2016) for each MiSeq run. First, the primer sequence, random hexamers, low quality and unexpectedly short reads were removed (parameters: truncLen = 240, 220, maxN = 0, maxEE = 2,2 for both group; trimLeft = 6 + 22, 6 + 25 for group 1, 6 + 29, 6 + 27 for group 2). Next, the error model was trained and used to identify and correct indel-mutations and substitutions of dereplicated passed sequences. The paired reads were then merged, and an amplicon sequence variant (ASV)–sample matrix was developed. To obtain a sample-specific standard line, linear regression analysis was performed using the read number of internal standard DNAs and their known copy numbers (the intercept was set at zero; lm function in R version 4.1.1 software; (R Core Team, 2021). For each sample, the number of eDNA copies for haplotypes was calculated using the sample-specific standard line: number of eDNA copies = the number of reads/regression slope of the sample-specific standard line (Table S3).

The species assignment for unique merged sequences was performed using a local BLASTN search with the reference database developed using the makeblastdb function (Camacho et al., 2009). The reference database for each group consisted of sequences downloaded from NCBI and standard DNA sequences for each group (Fig. S2a, b). For each unique sequence, top BLAST hits with sequence identity ≥ 96% were assigned as a species name. Only assigned sequences for each species (i.e. haplotypes) and standard DNA were used in subsequent analyses. The identity percentage (≥ 96%) used in the local BLASTN search was set slightly above the sequence similarity between the most closely related species (95.3%, between the W. Kyushu group of *O. obscura* and *O. hikimius*) to correctly assign species and detect as many haplotypes of each species as possible. To test the effects of this setting, we re-analysed the data using ≥ 98% identity, and the number of detected haplotypes and phylogenetic trees were compared with those obtained using ≥ 96% identity percentage (see below).

### 2.7 Data screening and phylogenetic analyses

DADA2 can remove most erroneous sequences generated during PCR and sequencing from raw sequence data, but some erroneous sequences may remain in the data without being removed (Tsuji et al., 2020a, 2020b). Data screening is key to eliminate remaining erroneous sequences and allow accurate haplotype detection based on eDNA analysis, as they produce noise in phylogenetic and phylogeographic analyses. We developed a data screening procedure to eliminate false positive sequences and improve the detection reliability of genetic lineages (an overview of the data screening flow was shown in Fig. S3). Our proposed data screening procedure primarily aimed at extracting certain major haplotypes in each site by removing error sequences as much as possible; accordingly, we boldly screened and removed haplotypes. Data screening was performed by species, but *O. obscura* and *O. hikimius* were analysed simultaneously as it is known that *O. obscura* does not form a monophyletic group (Fig. S2b).

The screening procedure of examined data consisted of the following three sequential procedures, as the additional effects can be assessed. First, haplotypes for which the estimated concentration was <1 copy/L filtered water volume were removed from the data (<1 copy/L replaced by 0; step 1), because such low concentration is theoretically unlikely. Second, haplotypes with a very low frequency (<1%) in each sample were removed from the data (<1% replaced by 0%; step 2), as they were suspected to be erroneous sequences and, even if real, they are unlikely to affect the result. Third, haplotypes with less than half or one-third of the proportion of the most predominant haplotype were removed from the data (<max%/2 replaced by 0%, step 3_1/2; or <max%/3 replaced by 0%; step 3_1/3). This assumes that individuals with the haplotypes characterising each regional lineage are likely dominant in each site, so their proportion of DNA copies will also be relatively high in each sample. Given the unknown appropriate frequency threshold, the less than half or one-third criterion was used to empirically determine a more suitable threshold for data screening based on reference sequence concordance and recovery rates (see below).

To examine the effectiveness of data screening, a phylogenetic tree was constructed for the obtained haplotypes in each step of the data screening procedure. The tree was estimated by the neighbour-joining (NJ) method (Saitou and Nei, 1987) using MEGA7 (Kumar et al., 2016) with the Jukes-Cantor model (Jukes and Cantor, 1969). Nodal support of the tree was assessed by bootstrap with 1,000 resamplings. In addition, when substantial sequence data are available as reference sequences, the proportion of haplotypes matching 100% with any reference sequences (100%-matched haplotypes) will increase if the haplotypes really exist. Therefore, we also used the number and proportion (%) of 100%-matched haplotypes as a performance indicator in each screening step. This examination was conducted only for *O. obscura* + *O. hikimius, N. temminckii* and *Z. platypus*; but not for *N. sieboldii*, because there were no 100%-matched haplotypes due to the lack of sufficient reference sequences (only one sequence available). To test whether we could exclusively exclude the haplotypes more likely to be false positives, the relationship between the decrease rate of a number of haplotypes by data screening and the proportion of 100%-matched haplotypes was examined by the generalised linear mixed model (GLMM) with a gamma distribution implemented by the ‘glmer’ function in the ‘lme4’ package ver.1.1-29 (Bates et al., 2022) for R, with the significance level set at α = 0.05. In this model, the proportion of 100%-matched haplotypes (matching rate) was set as a response variable, the decrease rate in the number of haplotypes detected in each step compared to that before data screening (step 0) as explanatory variable and the target species as random effects. Additionally, to examine the risk of removing real haplotypes (i.e. existing haplotypes in a sample) by data screening, the recovery rate of known haplotypes was calculated at each screening step.

Geographical distribution maps of genetic lineages were generated for the haplotype data obtained in the screening step 3 (<max%/2 replaced by 0%), which yielded the highest matching rate with the reference sequences (see Results and Table 1), using the R package ‘maps’ ver. 3.4.0 (Becker et al., 2021), ‘mapdata’ ver. 2.3.0 (Brownrigg, 2018) and ‘mapplots’ ver. 1.5.1 (Gerritsen, 2018). To examine how accurately the geographical distribution patterns of genetic lineages obtained by eDNA analysis reflect those obtained by conventional Sanger sequencing-based method, we directly compared the topology of NJ trees and distribution maps between the two methods.

**Table 1.** Summary of the number of haplotypes obtained in each step of the data screening process and those exactly matching any of the reference sequences (≥ 96% identity percentage). Data screening step: 0, non-data screening; 1, <1 copy/L replaced with 0 copy/L; 2, <1% in frequency at each site replaced with 0%; 3_1/2, the haplotypes detected at less than half of the proportion of the most predominant haplotype replaced by 0%; 3_1/3, the haplotypes detected at less than half of the proportion of the most predominant haplotype replaced by 0% (see Fig. S3)

| Target species | Data screening step | Total detected no. of haplotypes | Decrease rate from step 0 (%) | Number of haplotypes 100% matching with references | 100% Matching rate (%) | Recovery rate of references (%) | Figure of results |
| --- | --- | --- | --- | --- | --- | --- | --- |
| *Odontobutis obscura* and *O. hikimius*<br>No. of reference haplotypes: 5 | step 0 | 102 | 0.0 | 3 | 2.9 | 60.0 | Fig. S5a |
|  | step 1 | 92 | 9.8 | 3 | 3.3 | 60.0 | Fig. S5b |
|  | step 2 | 54 | 47.1 | 3 | 5.6 | 60.0 | Fig. S5c |
|  | step 3_1/2 | 19 | 81.4 | 3 | 15.8 | 60.0 | Fig. 2 |
|  | step 3_1/3 | 20 | 80.4 | 3 | 15.0 | 60.0 | Fig. S5d |
| *Nipponocypris temminckii*<br>No. of reference haplotypes: 3 | step 0 | 269 | 0.0 | 3 | 1.1 | 100.0 | Fig. S6a |
|  | step 1 | 163 | 39.4 | 3 | 1.8 | 100.0 | Fig. S6b |
|  | step 2 | 89 | 66.9 | 3 | 3.4 | 100.0 | Fig. S6c |
|  | step 3_1/2 | 18 | 93.3 | 3 | 16.7 | 100.0 | Fig. 3 |
|  | step 3_1/3 | 18 | 93.3 | 3 | 16.7 | 100.0 | Fig. S6d |
| *Zacco platypus*<br>No. of reference haplotypes: 33 | step 0 | 747 | 0.0 | 17 | 2.3 | 51.5 | Fig. S7a |
|  | step 1 | 517 | 30.8 | 17 | 3.3 | 51.5 | Fig. S7b |
|  | step 2 | 148 | 80.2 | 15 | 10.1 | 45.5 | Fig. S7c |
|  | step 3_1/2 | 39 | 94.8 | 12 | 30.8 | 36.4 | Fig. 4 |
|  | step 3_1/3 | 46 | 93.8 | 12 | 26.1 | 36.4 | Fig. S7d |

### 2.8 Sanger sequencing of tissue DNA

In addition to sequence data from previous studies, we newly determined sequences for several specimens of *O. obscura* and *N. sieboldii* by Sanger sequencing. In *O. obscura*, we newly detected haplotypes belonging to an unreported lineage group (E. Kyushu group, see Results) by eDNA. To confirm the presence of such haplotypes, we sequenced the partial mtDNA 12S rRNA gene (777 bp) for 10 individuals caught in sts. 88 and 89 (the Fukuchi River, Fukuoka Prefecture, Kyushu). A total of 110 *N. sieboldii* specimens were collected from 15 sites in western Japan (Table S4) and the mtDNA cyt*b* gene was sequenced. Note that the sequencing target was not the D-loop used for eDNA analysis, as it was performed for other purposes before the start of this study.

Total genomic DNA was isolated from a piece of fin or muscle tissue using a Wizard Genomic DNA Purification kit (Promega, Tokyo, Japan) or DNeasy blood and tissue kit. The 12S rRNA region of *O. obscura* (777 bp) and cyt*b* gene region of *N. sieboldii* (1237 bp, including flanking regions) were amplified using the following primer pairs, respectively: Odontobutis_sanger-F (5′-AGG GCC AGT AAA ACT CGT GC-3′) and Odontobutis_sanger-R (5′-GGG CGT CTT CTC GGT GTA AG-3′) (manually developed in this study), and L14724 (5′-TGA CTT GAA RAA CCA YCG YYG-3′) (Palumbi et al., 1991) and H15915 (5′-ACC TCC GAT CTY CGG ATT ACA AGA C-3′) (Aoyama et al., 2000). For *N. sieboldii*, only the 3′-half of the amplified region was sequenced using H15915 (715 bp). See the Appendix for details on PCR temperature conditions, product purification and Sanger sequencing. All sequences obtained were deposited in the International Nucleotide Sequence Database Collaboration database (accession No. LC719969–LC719978 for *O. obscura*; LC718524– LC718549 for *N. sieboldii*).

## 3. Results

### 3.1 Amplification tests of designed group primer sets

The *in silico* specificity check for each designed group-primer set by Primer-BLAST indicated group-specific amplification for the Odon primers. The NipZac primers would amplify seven *Cyprinus carpio* sequences (captured in China; NCBI accession No. FJ655351–FJ655357) as well as three target species of group 2 and *Opsariichthys uncirostris*. The *in vitro* amplification check for the Odon primers showed clear bands for the target DNA of *Odontobutis obscura* and *O. hikimius*. For the NipZac primer test, *C. carpio* DNA (captured at Lake Biwa, Japan; introduced Eurasian strain) was used in addition to the three target species and *O. uncirostris*. As a result, DNA amplification was found for only three target species (*N. temminckii, N. sieboldii* and *Z. platypus*) as well as for *O. uncirostris*.

### 3.2 Estimation of DNA copy number

The sequence reads of the internal standard DNAs were detected in all field-collected samples. For each primer set, sample-specific standard lines were successfully obtained by linear regression analysis (Table S3, Fig. S4). The slopes of sample-specific standard lines were highly variable, ranging from 33.0 to 1442.6 for the Odon primer set and from 11.7 to 2690.2 for the NipZac primer set. The *R*^2^ values of the regression lines were ≥ 0.96 except for one sample (st. 81) with 0.92 from the Odon primer set. The sequence reads of the unique sequences obtained with each primer set per sample were successfully converted to DNA copy number using the regression slope of the obtained sample-specific standard line.

### 3.3 Sequence assignment to species and data screening

The local BLASTN search with a 96% identity percentage successfully assigned unique sequences to target species and standard DNAs (Table S5). Additionally, unmatched sequences (i.e. sequence identity <96%) were also detected in all MiSeq runs. Several haplotypes were shared between different MiSeq runs; the total number of haplotypes for each target species finally obtained from the four MiSeq runs (before screening) was as follows: *O. obscura* 102, *O. hikimius* 17, *N. temminckii* 269, *N. sieboldii* 15 and *Z. platypus* 747 haplotypes (Table S6).

Data screening was performed in three steps (Fig. S3). The total number of detected haplotypes and the obtained NJ trees in each data screening step are shown in Table 1 and Figs. S5, S6 and S7. For all four species for which screening was applicable (*O. obscura, O. hikimius, N. temminckii* and *Z. platypus*), the number of haplotypes detected decreased with increasing data screening step; the decrease rate being greatest between steps 2 and 3. Additionally, the matching rate with reference sequences was significantly positively related to the haplotype decrease rate from step 0 (i.e. non-data screening) (*p* < 0.001, GLMM; Fig. S8). For *O. obscura* + *O. hikimius* and *Z. platypus*, the matching rate was greatest in step 3_1/2 For *N. temminckii*, the matching rate was a tie for the highest at steps 3_1/2 and 3_1/3 (Table 1). The decrease rate in the number of haplotypes from step 0 ranged between 80%–90% in steps 3_1/2 and 3_1/3. For *O. obscura* + *O. hikimius* and *N. temminckii*, the recovery rate of the reference haplotypes did not change with or without data screening or through its progression (60% and 100%, respectively); however, for *O. platypus*, the recovery rate decreased by 15% from step 0 (51.5%) to step 3 (36.4%) (Table 1).

When reanalysed with a ≥ 98% identity percentage for the local BLASTN search, the same or slightly fewer haplotypes were detected by data screening steps 3_1/2 and 3_1/3 compared to the results using ≥ 96% identity percentage (Figs. S9, S10, S11, S12). For *O. obscura* + *O. hikimius, N. sieboldii* and *Z. platypus*, regardless of the frequency threshold used in data screening step 3, the NJ tree for the haplotypes detected with ≥ 98% identity was similar to that obtained when the BLASTN search was performed using ≥ 96% identity. However, for *N. temminckii*, group C (Tokai region) was not detected at all at ≥ 98% identity.

### 3.4 Comparison of phylogeographic patterns between eDNA and conventional methods

For all target species, the major genetic lineage groups and their geographic distribution estimated using eDNA analysis were in almost perfect agreement with those obtained in Sanger sequencing-based studies (Figs. 2, 3, 4, 5). Furthermore, haplotypes belonging to subgroups with relatively high bootstrap values within each lineage group had a regionally restricted distribution.

**Fig. 2.**
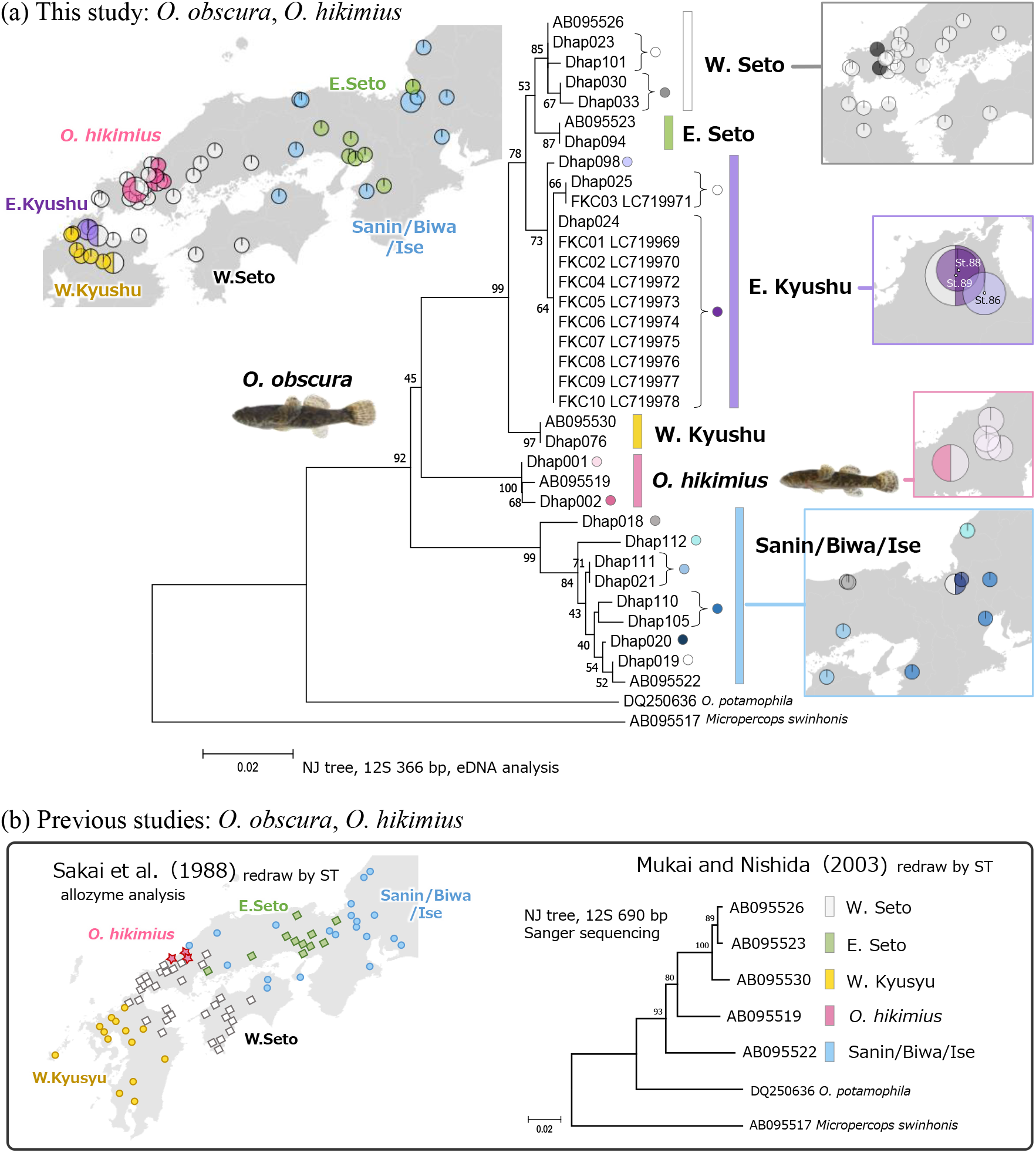
*Odontobutis obscura* and *O. hikimius*; (a) NJ tree and distribution map based on partial 12S sequence (366 bp) obtained by eDNA analysis and (b) NJ tree based on the deposited partial 12S sequence (690 bp) in NCBI by Mukai and Nishida (2003) and distribution map of each group revealed using allozyme analysis by Sakai et al. (1988). Numbers at internodes of both NJ trees represent bootstrap probability values (≥ 40 %) for 1,000 replicates. The colours of each group are common in both panels, NJ trees and distribution maps. IDs in NJ trees: ‘Dhap No.’, haplotype detected by eDNA analysis; ‘FKC No.+ LC7199xx’, ID and accession No. of individuals captured and sequenced in st. 88 and 89 (Fukuchi River); ‘AB0955xx.’, accession No. of deposited sequence in NCBI by Mukai and Nishida (2003). Pie chart shows the ratio of detected haplotypes of each group and the relative total number of haplotypes detected (Table S7a).

**Fig. 3.**
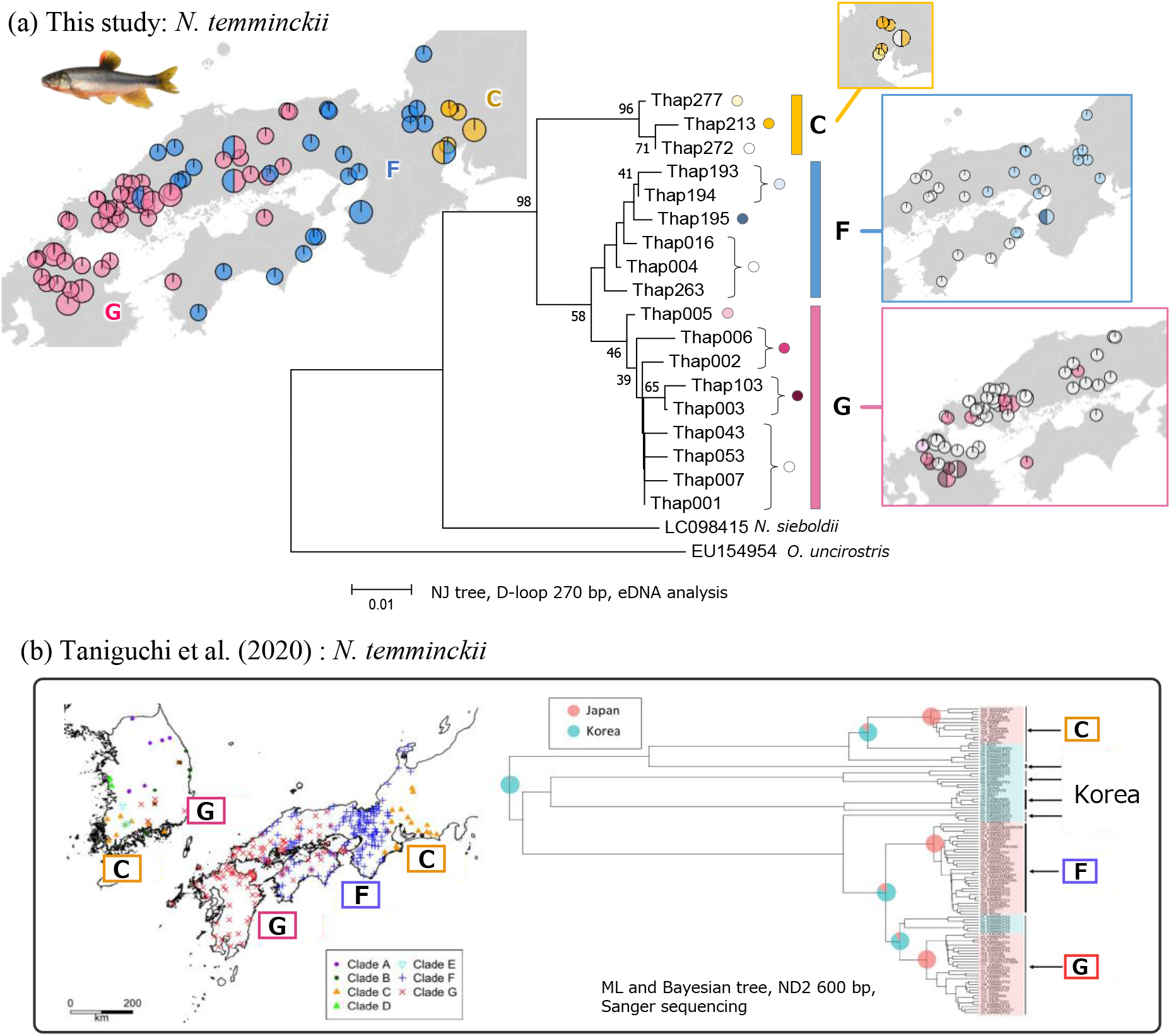
*Nipponocypris temminckii*; (a) NJ tree and distribution map based on partial D-loop sequence (270 bp) obtained by eDNA analysis and (b) ML and Bayesian tree and distribution map based on partial ND2 sequence (600 bp) provided by Taniguchi et al. (2020). Numbers at internodes of NJ tree represent bootstrap probability values (≥ 30 %) for 1,000 replicates. The colours of each group are common in the panels, trees and distribution maps. IDs in NJ trees: ‘Thap No.’, haplotype detected by eDNA analysis. Pie chart shows the ratio of detected haplotypes of each group and the relative total number of haplotypes detected (Table S7b).

**Fig. 4.**
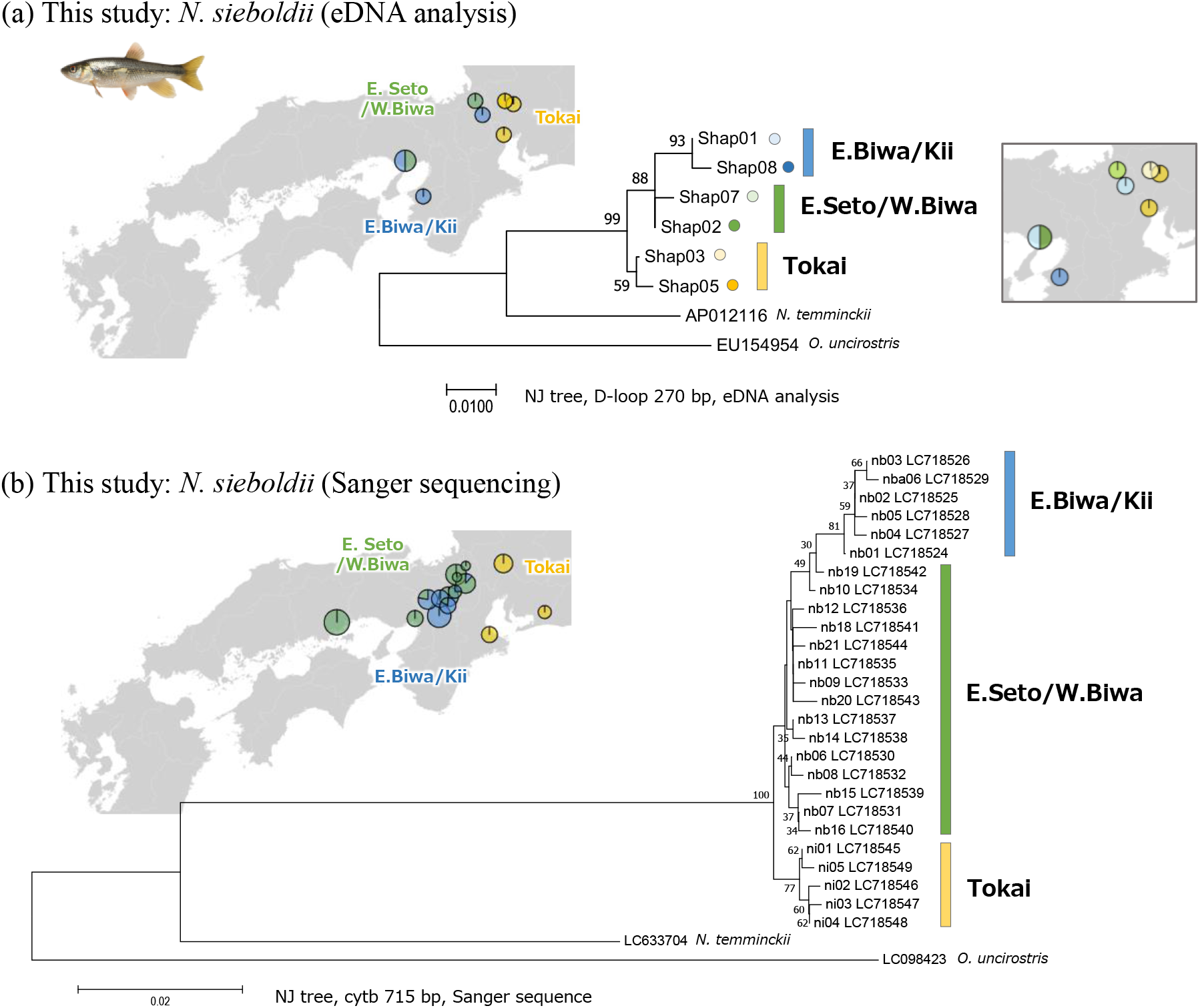
*Nipponocypris sieboldii*; NJ tree and distribution map (a) based on partial D-loop sequence (270 bp) obtained by eDNA analysis and (b) based on partial cytb sequence (715 bp) obtained by Sanger sequence. Numbers at internodes of both NJ trees represent bootstrap probability values (≥ 30 %) for 1,000 replicates. The colours of each group are common in the panels, NJ trees and distribution maps. IDs in NJ trees: ‘Shap No.’, haplotype detected by eDNA analysis; ‘LC7185xx’, NCBI accession No. Pie chart shows the ratio of detected haplotypes of each group and the relative total number of haplotypes detected (Table S7c).

**Fig. 5.**
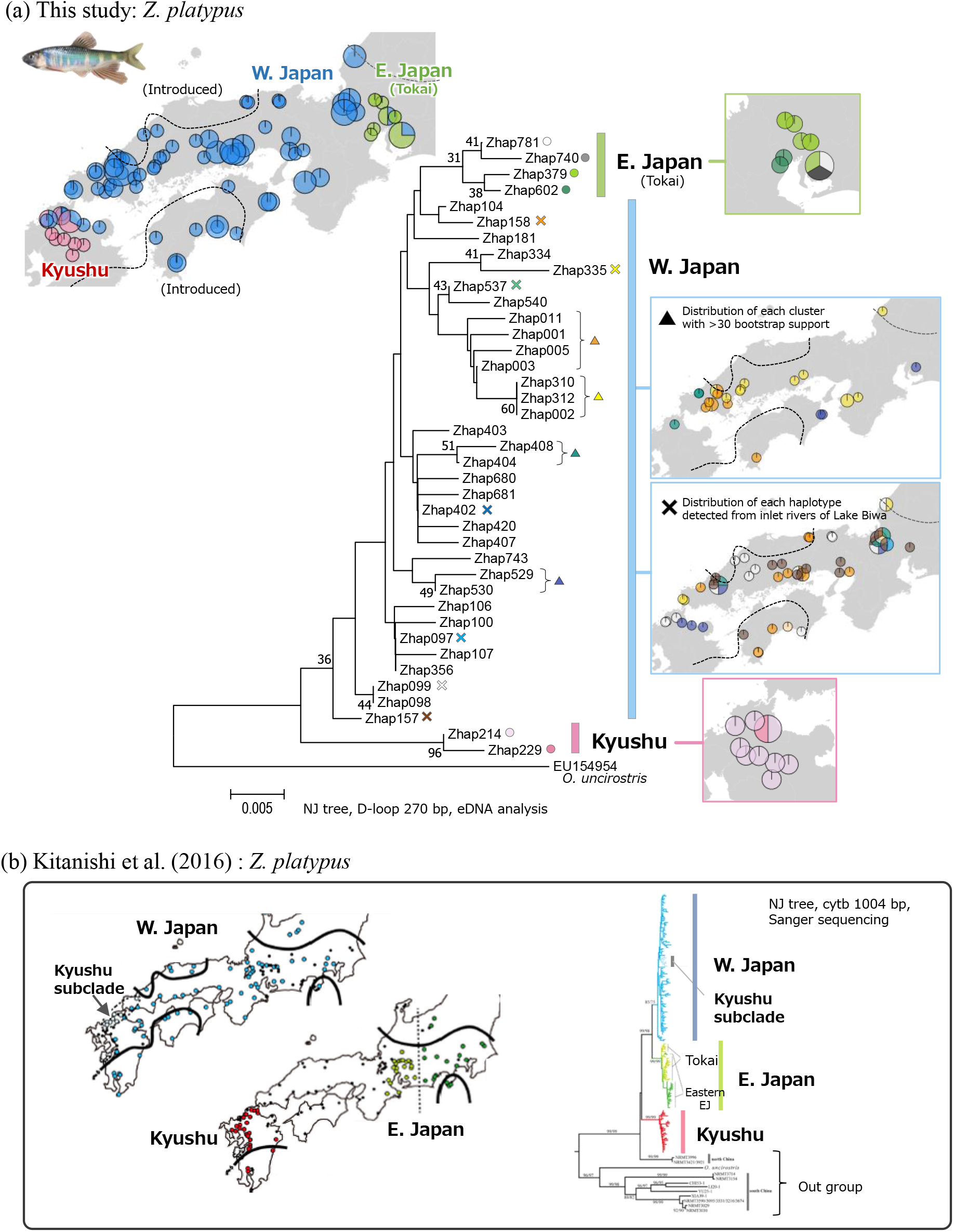
*Zacco platypus*; NJ tree and distribution map (a) based on partial D-loop sequence (270 bp) obtained by eDNA analysis and (b) based on partial cytb sequence (1,004 bp) provided by Kitanishi et al. (2016). Numbers at internodes of NJ tree in (a) represent bootstrap probability values (≥ 30 %) for 1,000 replicates. The colours of each group are common in the panels, NJ trees and distribution maps. IDs in NJ trees: ‘Zhap No.’, haplotype detected by eDNA analysis. Pie chart shows the ratio of detected haplotypes of each group and the relative total number of haplotypes detected (Table S7d).

#### 3.4.1 Odontobutisobscura and O. hikimius

A total of 17 and two haplotypes of *O. obscura* and *O. hikimius* were detected from a total of 49 and five study sites, respectively (Fig. 2, Table S2). Of a total of 19 haplotypes, three were known haplotypes (recovery rate 60% in all data screening steps) and the remaining 16 were newly detected (Table 1). In addition to the four lineage groups shown in previous studies (Sakai et al., 1988; Mukai and Nishida, 2003), the East Kyushu group was newly found by eDNA analysis in sts. 88, 89 (Fukuchi River) and 86 (Ima River) (Table S7a). A later capturing survey and Sanger sequencing confirmed that all *O. obscura* specimens caught in the Fukuchi River had haplotypes belonging to the East Kyushu group, which were in perfect agreement with those detected from the Fukuchi River in eDNA analysis (haplotype ID, Dhap024 or Dhap025; Fig. 2). Furthermore, several subgroups with clear regional specificity within the West Seto and Sanin/Biwa/Ise groups of *O. obscura* were revealed.

#### 3.4.2 Nipponocypris temminckii, N. sieboldii and Zacco platypus

A total of 18, 6 and 39 haplotypes of *N. temminckii, N. sieboldii* and *Z. platypus* were detected from a total of 79, 8 and 78 study sites, respectively (Table S2, Figs. 3, 4, 5). Of all detected haplotypes for *N. temminckii* and *Z. platypus*, three (recovery rate 100% in all data screening steps) and 12 (final recovery rate 36.4% in data screening step 3_1/2) were known haplotypes and the remaining 15 and 27 were newly detected (Table 1). In each of the three species, as indicated by Sanger sequencing based studies (Taniguchi et al., 2021, this study and Kitanishi et al., 2016), three major lineage groups were identified. Of those, a genetic group unique to the Tokai region was found in common among the three species (C group for *N. temminckii*; Tokai group for *N. sieboldii*; E. Japan group for *Z. platypus*). For *Z. platypus*, several haplotypes detected in the inlet rivers of Lake Biwa (the western Japan group) were found throughout the study area (Fig. 5).

## 4. Discussion

### 4.1 Group-specific primer development

The results of the specificity tests consistently showed that the developed two group-specific primers was sufficiently specific for this study. With respect to the Odon primer set, the results of both *in silico* and *in vitro* tests suggested that the primers can specifically amplify the DNA of *Odontobutis obscura* and *O. hikimius*. For the NipZac primer set, an *in silico* test could amplify the *Cyprinus carpio* DNA, but *in vitro* tests using tissue DNA from *C. carpio* caught in Japan (introduced Eurasian strains) confirmed that they were not amplified. The risk of false amplification is not zero, but even if it occurs, it would not affect the results as it can be excluded during data analysis. Furthermore, both primer sets were group-specific, as none of the sequences detected with each primer from the field samples matched non-target species other than *O. uncirostris*, which was allowed to amplify.

### 4.2 Species assignment of detected sequences after denoising

Each group-primer set amplified the DNA of multiple target species, and each sequence detected could be correctly assigned to a target species by a local BLASTN with ≥ 96% identity. In this study, to avoid missing any existing genetic lineages in each species, we set the identity percentage as low as possible, i.e. slightly above the sequence identity between the most closely related species (95.3%). In eDNA-based phylogeography, a more conservative (i.e. higher) species identity percentage would increase the possibility of false negative detection of haplotypes from unknown, distant lineages. This risk was supported by the fact that the C group (Tokai region) of *N. temminckii* was not detected in the re-analysis with an identity threshold ≥ 98% (Fig. S10). Given the trade-off between accurate species assignment and comprehensive detection of intraspecific lineages, the identity percentage needs to be carefully determined as low as possible within which the target species can be distinguished.

### 4.3 Data screening for increased reliability of the results

Although previous studies demonstrated that denoising of FASTQ files using appropriate algorithms can remove erroneous sequences, completely removing them is still challenging in some cases (Callahan et al., 2016; Edgar, 2016; Tsuji et al., 2020b; Turon et al., 2020) In our all MiSeq run data, after the species assignment using the local BLAST, numerous standard DNA sequences containing a few SNP-level errors and chimeric sequences containing standard DNA-specific ones were also detected (Table S5). Based on these observations, it is reasonable to assume that such SNP-level errors and chimeric sequences were still contained in the assigned sequences to each target species.

Our proposed bold data screening procedure primarily aimed at removing error sequences as much as possible; there is a trade-off between data screening successs and the risk of accidentally removing real minor haplotypes. However, even if parts of real minor haplotypes are incorrectly removed, it would not affect much the estimation of major lineage groups and their geographical patterns, as relatively frequent haplotypes will remain in each study site. With progressing data screening, the number of detected haplotypes decreased, ultimately excluding ≥ 80% in step 3, but the matching rate to reference sequences significantly increased. Additionally, the recovery rate of the reference haplotypes changed little before and after data screening. These results suggest that the proposed data screening procedure selectively eliminated erroneous sequences (false positives) that could not be removed by denoising using DADA2, moderately reducing the occurrence of false negatives.

To determine the appropriate frequency threshold for data screening step 3, we examined the threshold based on the matching rate with and recovery rate of reference sequences. For all species considered, the recovery rates did not differ between the two thresholds (1/2 vs. 1/3), but the matching rate was slightly higher at the 1/2 threshold for *O. obscura* + *O. hikimius* and *Z. platypus*. Thus, we used 1/2 as threshold for data screening step 3 in our subsequent analyses. However, further studies are necessary for the frequency threshold because the risk of false-negative results positively correlates with the threshold value.

### 4.4 Usefulness and potential of eDNA-based phylogeography

The phylogenetic trees and geographical distribution patterns of genetic lineage groups estimated using eDNA almost perfectly reflect the results obtained by Sanger sequencing, demonstrating the high usefulness and potential of eDNA analysis in phylogeography (Figs. 2, 3, 4, 5). These three findings are particularly noteworthy: (1) discovery of a new regional population group, (2) comparative phylogenetic inference, i.e. detection of common regional population groups among species and (3) detection of artificial distribution disturbance (Fig. 5).

With respect to the first point, multisite surveys (total of 94 broadly-distributed sites), which were easily achieved due to the simplicity of the eDNA survey and reuse of previous samples, may have contributed to the discovery of the East Kyushu group of *O. obscura* (Fig. 2). This group was found only in two rivers (sts. 88 and 89, Fukuchi River and st. 86, Ima River), which previous studies failed to sample. In addition, several subgroups with clear regional specificity were revealed within major lineage groups (e.g. the Sanin/Biwa/Ise group). A positive correlation between the density of the survey sites and the detectability of local lineage groups is highly plausible, especially for species with high regional-specificity, such as *O. obscura*. Although it is preferable to sample from as many sites as possible, many site capture surveys covering the distribution range of the target species usually require an enormous amount of time and effort even when targeting a single species. Therefore, we propose conducting an exhaustive survey with cost-effective eDNA analysis, followed by intensive capture surveys at interesting sites. This survey strategy would increase the efficiency and comprehensiveness of the survey while prompting more detailed phylogeographic studies based on tissue DNA analysis from captured specimens.

Second, eDNA-based phylogeography allows examining multiple species simultaneously using the same samples, facilitating comparative studies for phylogeographic structure among co-distributed species. The Tokai region, common for all three species in group 2 (*N. temminckii, N. sieboldii* and *Z. platypus*), suggests that their population structure was affected by the geographical boundaries of the Ibuki–Suzuka Mountains (Fig. 1). The importance of this boundary has been documented in previous phylogeographic studies of several freshwater fish species (Miyazaki et al., 2011; Takehana et al., 2003; Tominaga et al., 2016; Watanabe et al., 2014). For comparative phylogeography based on capture surveys and Sanger sequencing, the cost of analysis increases with the number of species being compared. This practical problem is one of the major challenges in conducting comparative phylogeography. In this study, we designed group-primer sets for each of the two target groups containing two or three species and simultaneously determined the sequences of their amplicons in one library. The effects of the number of target species on the effort, time and cost needed for analysis are usually small, as only a single sequence library preparation and sequencing are required for eDNA-based comparative phylogeography. Therefore, the use of eDNA analysis has great potential to solve this cost problem, greatly acilitating comparative phylogeography.

Third, the distribution patterns of haplotypes revealed by eDNA analysis may also be helpful to monitor the invasion of non-native lineages. *Zacco platypus* is known to have been unintentionally introduced to almost all regions of Japan from Lake Biwa, mainly in association with the stocking of Ayu, *Plecoglossus altivelis*, one of the most important species for freshwater fisheries in Japan causing genetic disturbances (Kitanishi et al., 2016; Mizuguchi, 1990; Takamura and Nakahara, 2015). In our results, the haplotypes found in the inlet rivers of Lake Biwa were also detected throughout western Japan, suggesting that eDNA analysis could successfully reveal the current state of a genetic disturbance in *Z. platypus*. Anthropogenic species introduction is a serious problem for the conservation of freshwater ecosystems worldwide (Cucherousset and Olden, 2011; Gozlan et al., 2010). This is also the case in Japan, where many freshwater fish species have suffered genetic disturbance through fisheries stocking or arbitrary release by aquarium hobbyists (Miyake et al., 2011, 2021; Tominaga et al., 2020). However, such introductions, especially those between natural distribution areas, are ‘cryptic threats’, and it is difficult to ascertain their actual status without genetic analysis (Mukai et al., 2013). Since eDNA analysis is suitable for long-term monitoring (Rees et al., 2014; Székely et al., 2021), it would enable early recognition of invasive threats contributing to early conservation.

### 4.5 Limitations and future challenges of eDNA-based phylogeography

The high usefulness and potential of eDNA analysis in phylogeography are unquestionable, but thre are certain limitations and challenges. In particular, it is important to recognise the limitation of the length of DNA sequences that can currently be analysed in eDNA analysis. This limitation is largely related to the concentration and persistence of the target eDNA in the field. Recently, several studies have reported successful long-read sequencing from eDNA samples, suggesting the presence of nearly intact mitochondria and nuclei in water as a source for eDNA (Deiner et al., 2017; Jensen et al., 2021; Kakehashi et al., 2022). However, these studies were conducted with high eDNA concentrations of the target species (i.e. tank and high-density habitats); hence, in real-life conditions, those results would be hard to achieve. Indeed, in a study on the mackerel *Trachurus japonicus* in Maizuru Bay, Japan, false negatives or significant reductions in concentrations were observed at most survey sites by lengthening the target sequence length by 600 bp (from 127 to 719 bp) (Jo et al., 2017). Given these considerations, as the target sequence becomes longer, the risk of false negative results is likely to increase for species with smaller biomass and/or abundance.

On the other hand, the short sequences of mitochondrial DNA yield limited resolution results (Jensen et al., 2021). In this study, NipZac primers amplified 270 bp of the D-loop region of *Z. platypus*; we missed the Kyushu subclade shown in a previous study using 1,004-bp cyt*b* sequences (Kitanishi et al., 2016). This false negative result was most likely due to the insufficient resolution caused by the shortness of the analysed sequences. Thus, it is important to target as long sequences as possible to improve the detection of lineage groups. This may be achieved by conducting surveys during the spawning season when eDNA concentrations are temporarily much higher than usual due to released sperm (Bylemans et al., 2017; Tsuji and Shibata, 2021). This limitation of sampling time, however, sacrifices the ease of sampling.

If high concentrations of high-quality DNA can be recovered, it may also be possible to target the nuclear DNA, which has a lower copy number in the cell than mitochondrial DNA. The shortcomings of relying only on mitochondrial DNA to infer and discuss the population structure have long been recognised in many previous studies (Ballard and Whitlock, 2004; Teske et al., 2018). Future development of stable detection methods for nuclear DNA from eDNA samples will pave the way for the analysis of genomic variation in populations by environmental sampling, making eDNA-based phylogeography increasingly useful.

## 5. Conclusion

By comparing our results with known phylogeographic patterns for five freshwater fish species, this study demonstrated that eDNA analysis can be a useful tool for phylogeography. For all target species, the phylogenetic trees and geographical distribution patterns of genetic lineage groups estimated based on eDNA analysis through our proposed data screening procedure almost perfectly reflected those obtained by conventional methods using Sanger sequencing of tissue DNA. Despite some limitations and future challenges remain, the application of eDNA analysis to phylogeography can significantly reduce the time and effort of surveys and make multi-species targeted surveys much easier. The eDNA phylogeography will be increasingly studied in the future and will continue to grow into a more useful and powerful tool.

## Authors’ contributions

S.T., Y.A. and K.W. conceived and designed the research. S.T., N.S., R.I, R.N. and K.W. performed a field survey. S.T., N.S. and K.W. performed molecular experiments and data analysis. S.T. wrote the early draft and completed it with significant input from all authors.

## Data availability

All raw sequences were deposited in the DDBJ Sequence Read Archive (accession number: DRA014749).

## Acknowledgements

We sincerely thank laboratory members of Akamatsu laboratory, Yamaguchi University for helping in water sampling, T. Abe, A. Iwata, T. Shimizu, H. Yoshigo for providing *N. sieboldii* samples, and S. Kunimatsu for providing fish pictures. This study was supported by the Sasagawa Scientific Research Grant from the Japan Science Society (Grant No. 202-5001), ESPEC Foundation for Global Environment Research and Technology (Charitable Trust) and YU Project for Formation of the Core Research Center.

---

**Fig. S1.**
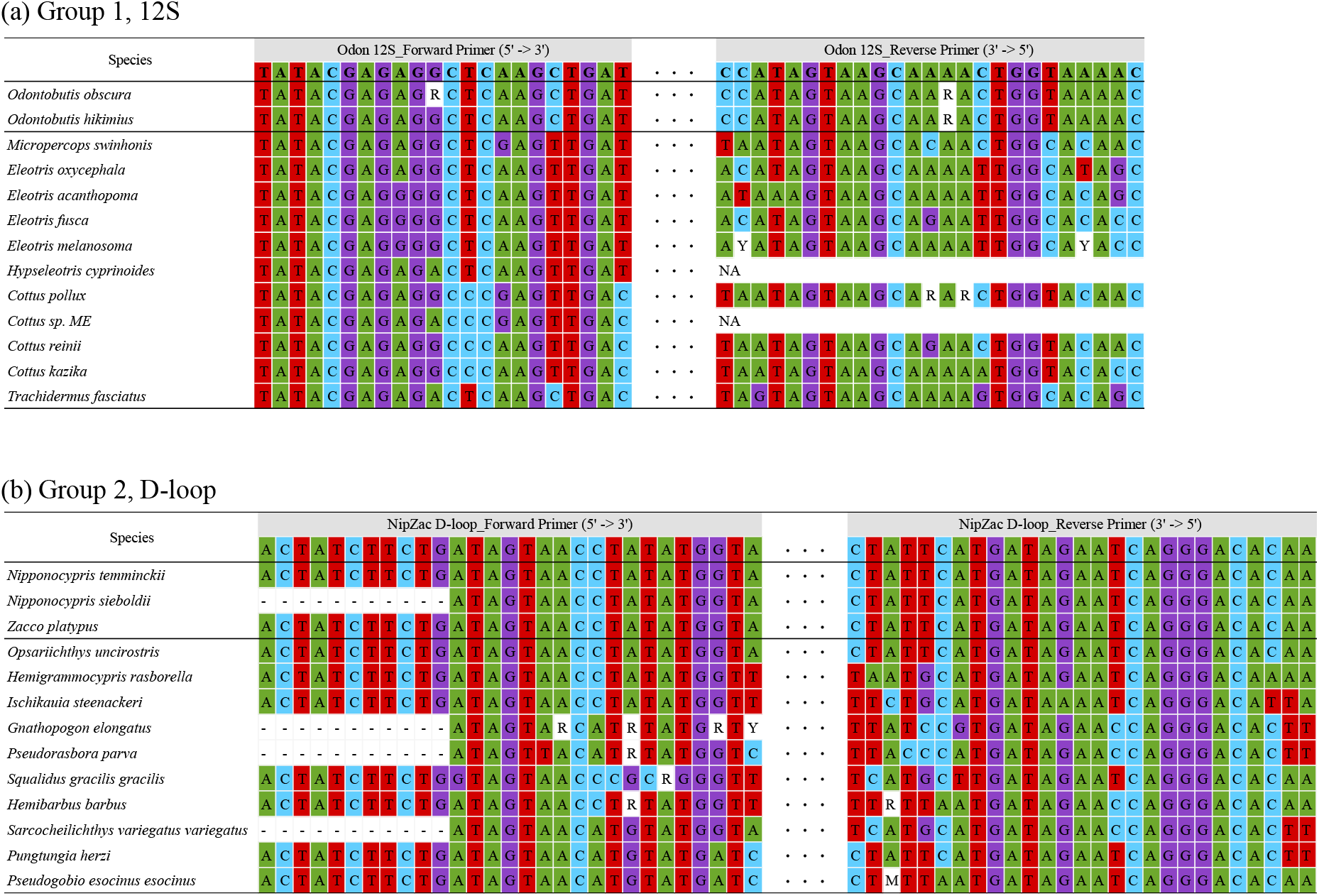
Primer specificity for (a) group 1 (*Odontobutis obscura* and *O. hikimius*) and (b) group 2 (*Nipponocypris temminckii, N. sieboldii* and *Zacco platypus*). The R, M and Y indicate (A or G), (A or C) and (C or T), respectively.

**Fig. S2a.**
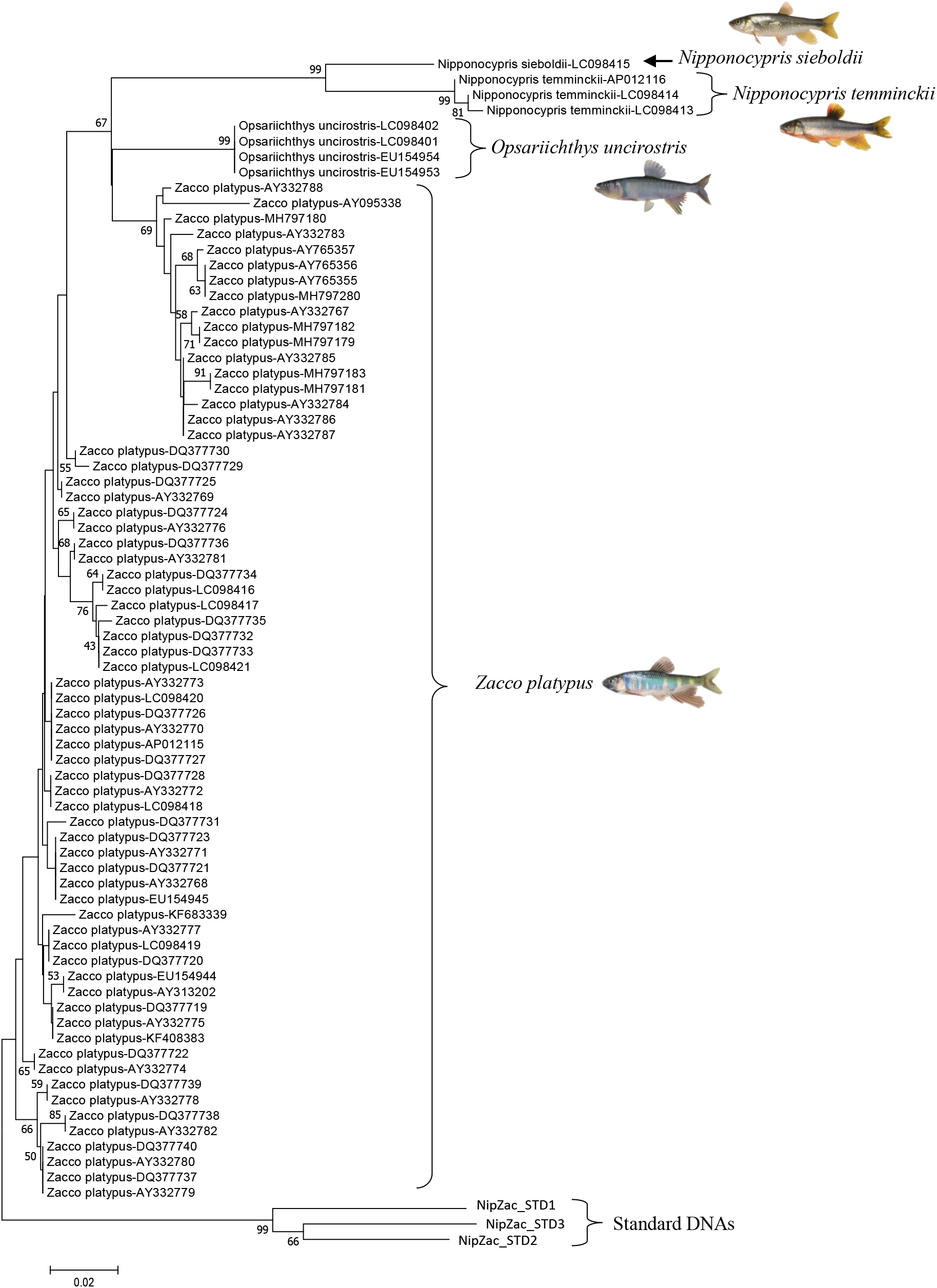
NJ tree of reference sequence used in species assignment step for NipZac primer. Numbers at internodes represent bootstrap probability values (≥ 40 %) for 1,000 replicates.

**Fig. S2b.**
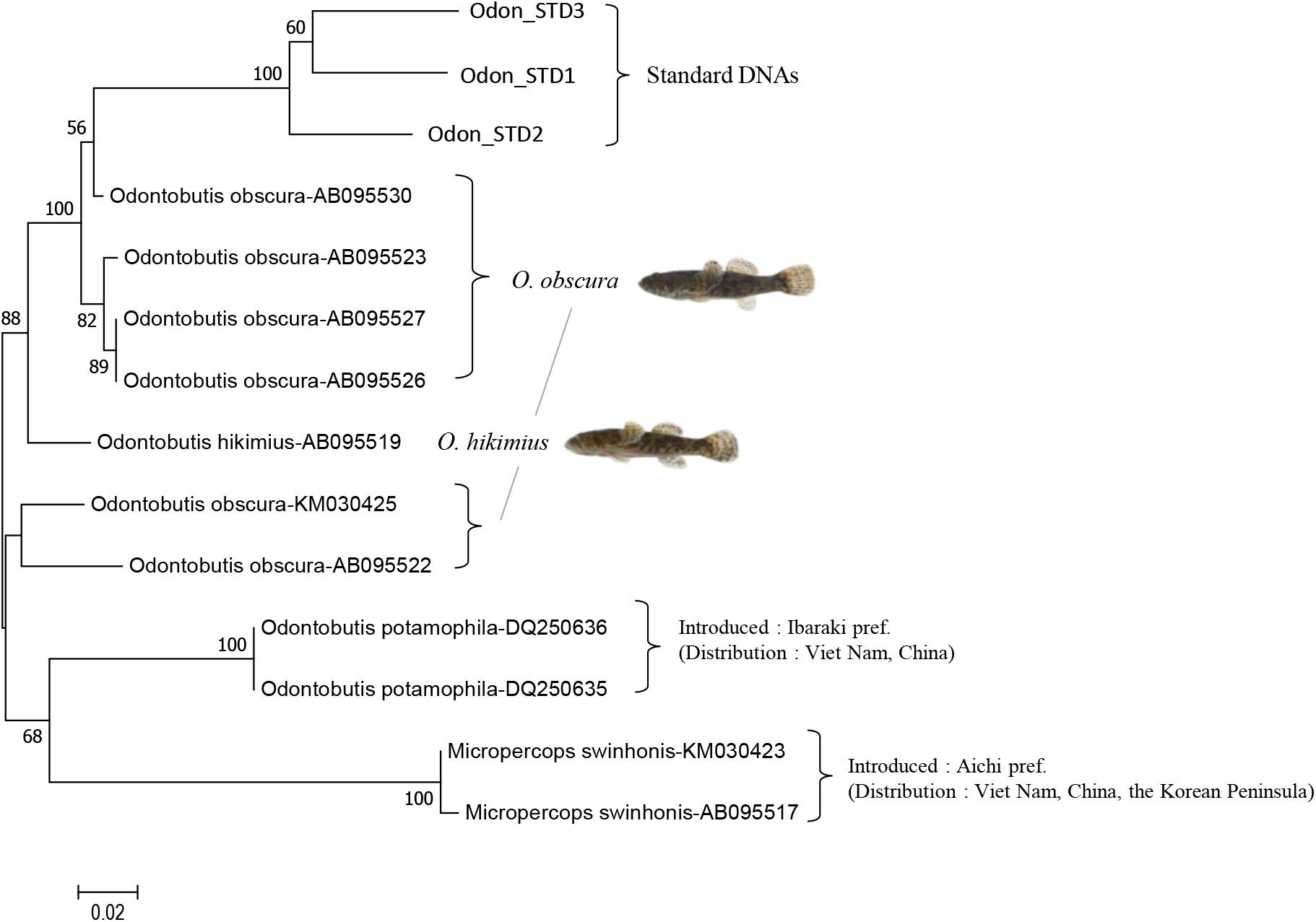
NJ tree of reference sequence used in species assignment step for Odon primer. Numbers at internodes represent bootstrap probability values (≥ 40 %) for 1,000 replicates.

**Fig. S3.**
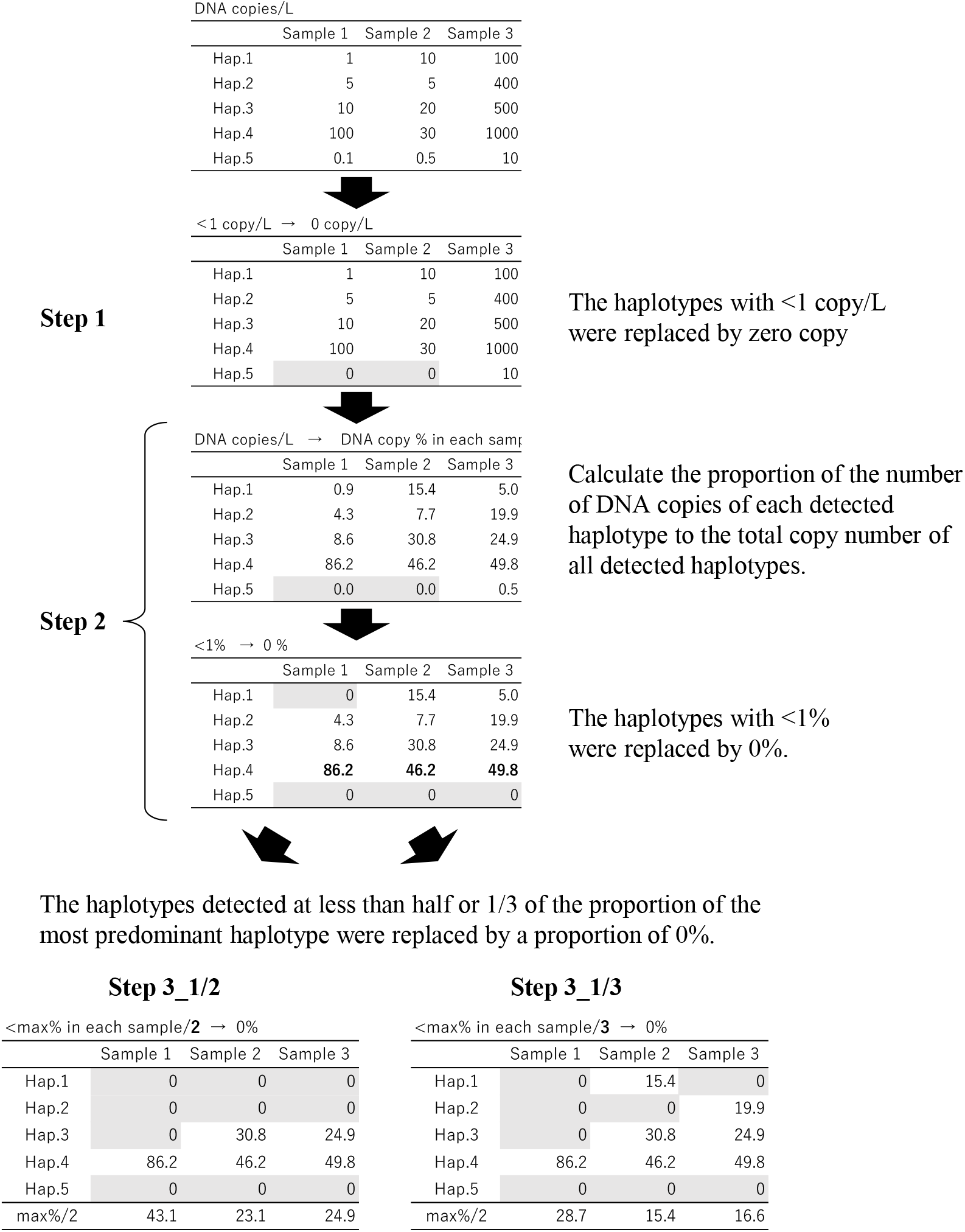
Overview of the data screening flow.

**Fig. S4.**
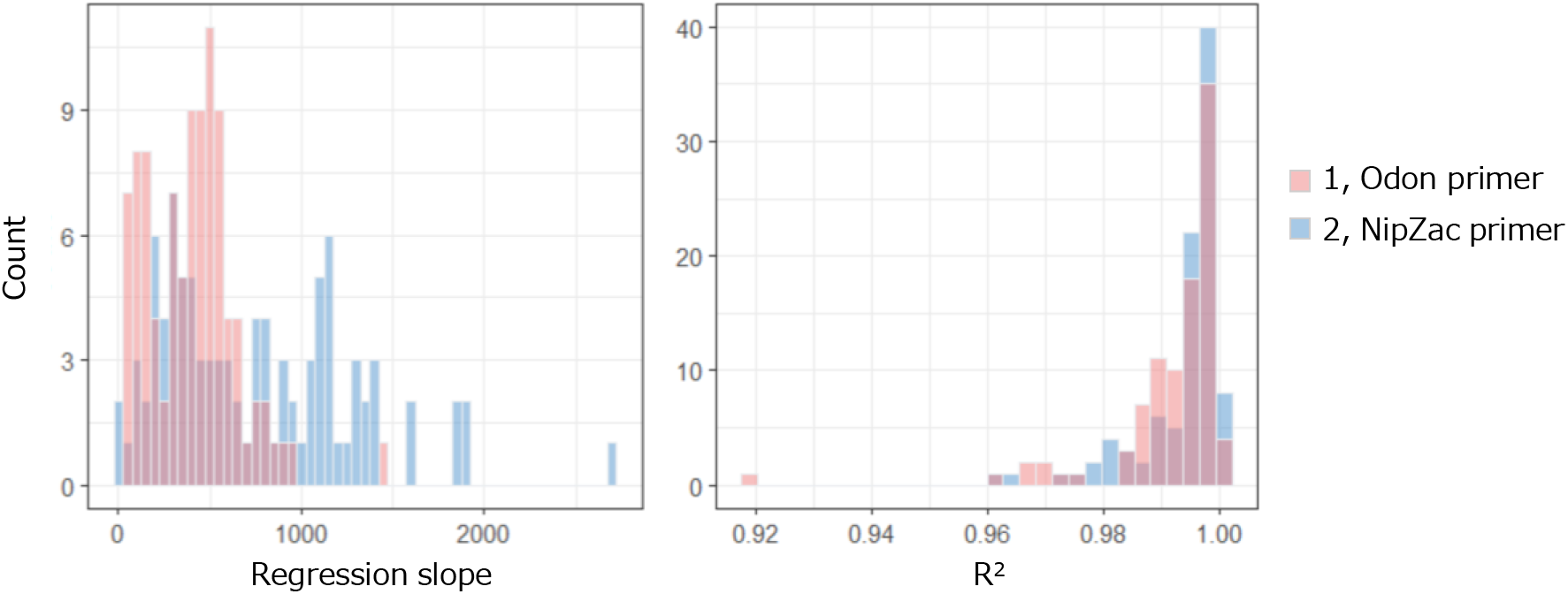
Summary of regression slope and *R*^2^ values of sample-specific standard lines constructed using the number of copies added and sequence reads of internal standard DNAs.

**Fig. S5a.**
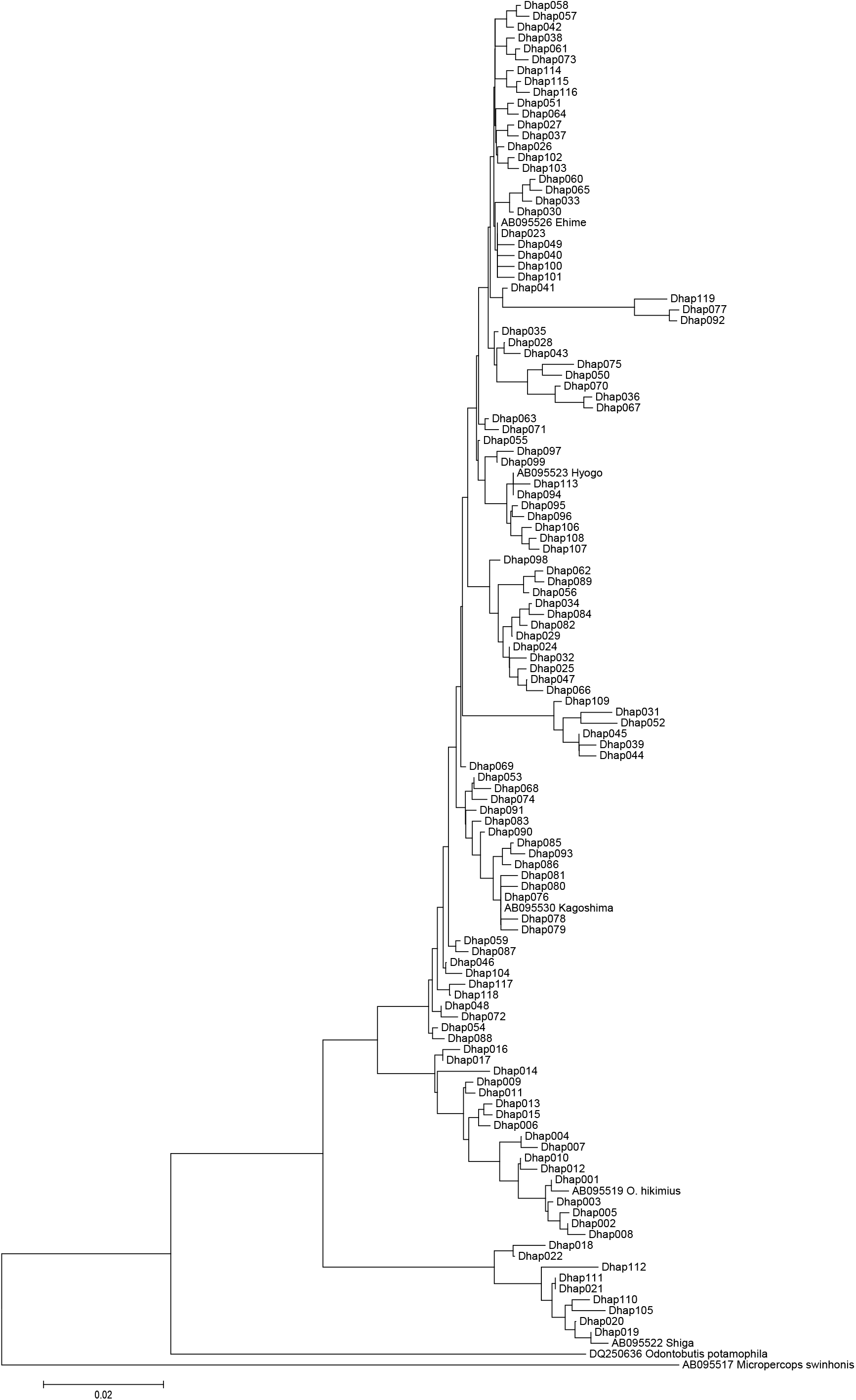
*O. obscura* and *O. hikimius:* NJ tree based on detected haplotypes (step 0, before data screening).

**Fig. S5b.**
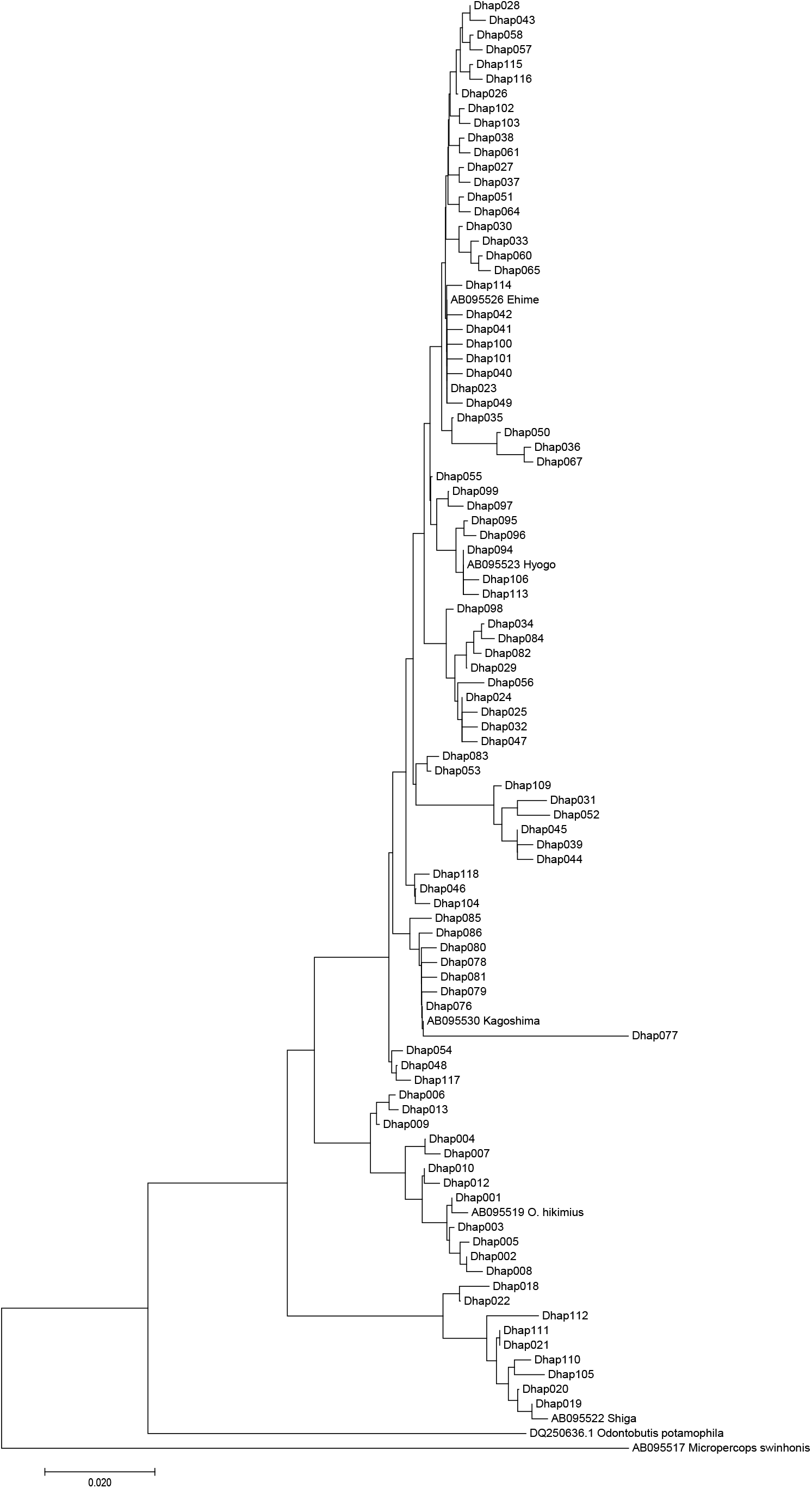
*O. obscura* and *O. hikimius:* NJ tree based on detected haplotypes after data screening step 1.

**Fig. S5c.**
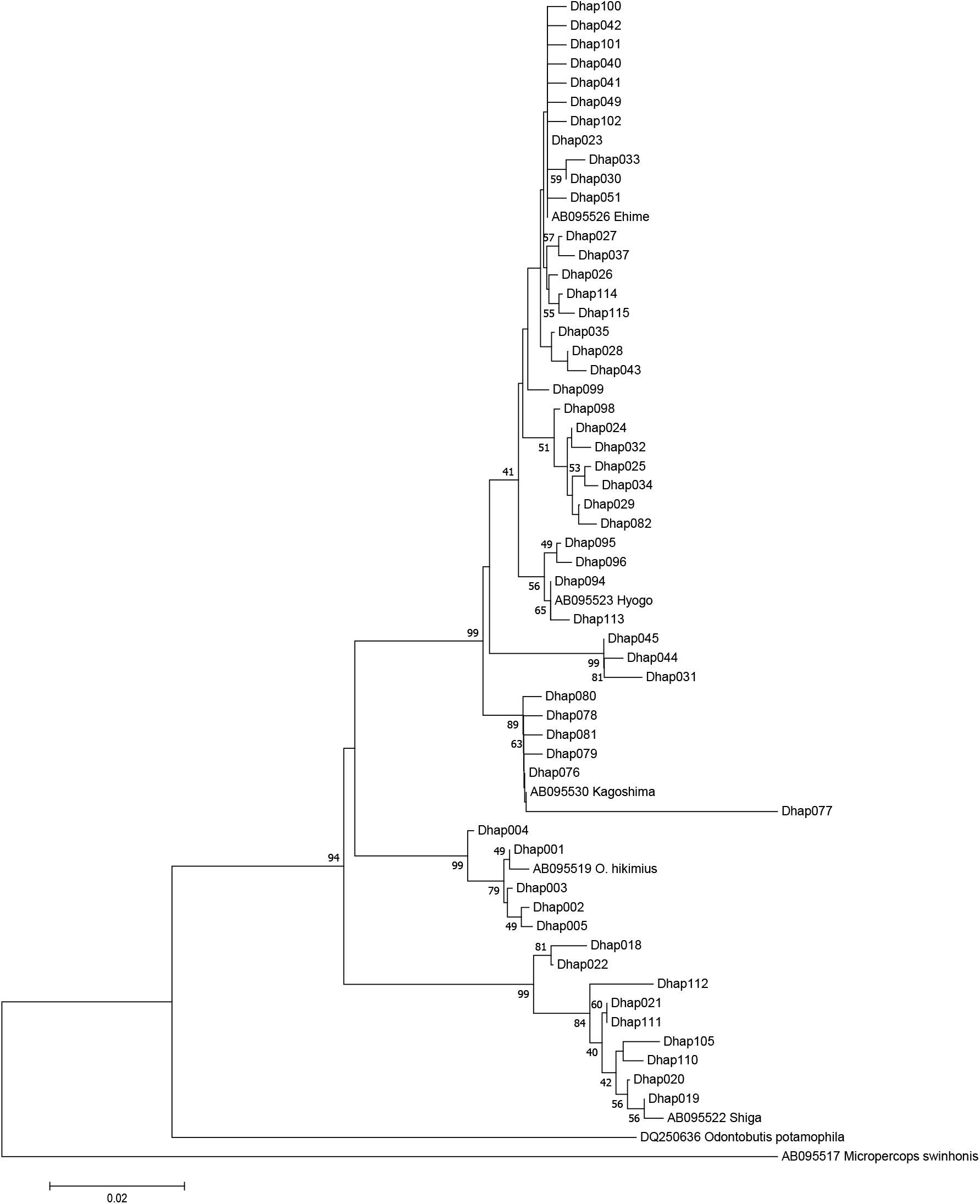
*O. obscura* and *O. hikimius:* NJ tree based on detected haplotypes after data screening step 2. Numbers at internodes represent bootstrap probability values (≥ 30 %) for 1,000 replicates.

**Fig. S5d.**
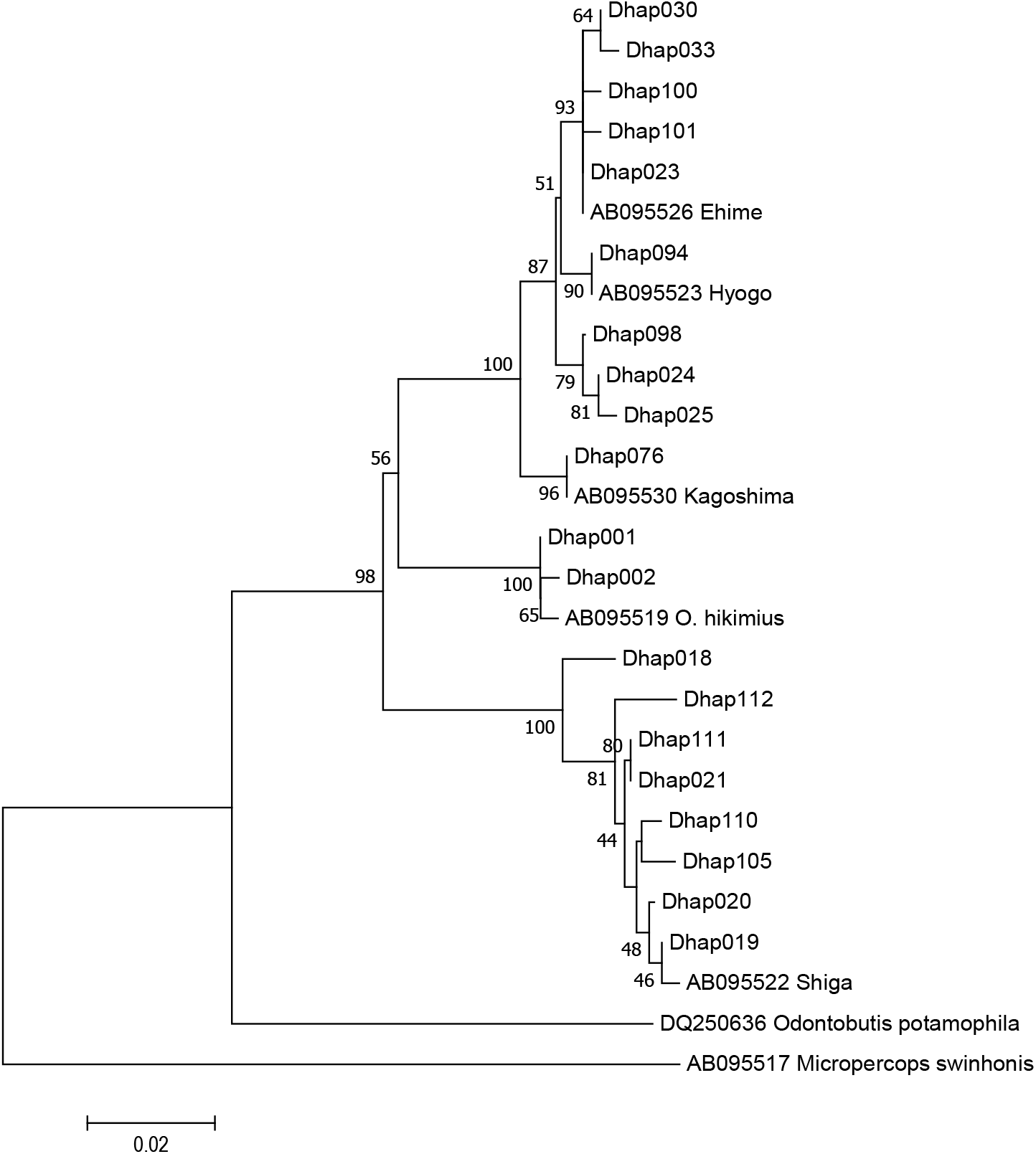
*O. obscura* and *O. hikimius:* NJ tree based on detected haplotypes after data screening step 3_1/3 (<max%/3 → 0%). Numbers at internodes represent bootstrap probability values (≥ 30 %) for 1,000 replicates.

**Fig. S6a.**
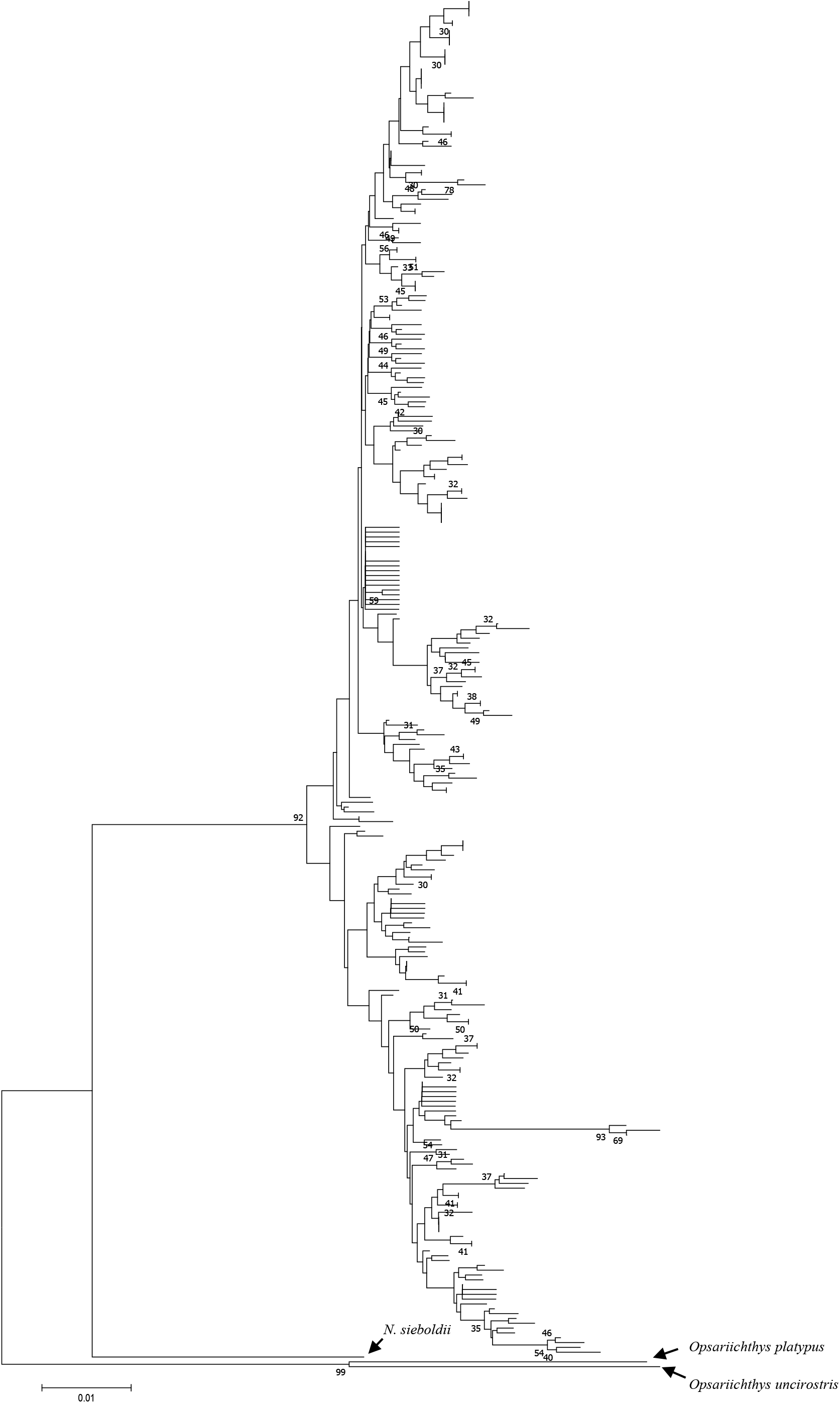
*N. temminckii*: NJ tree based on detected haplotypes (step 0, non-data screening). Numbers at internodes represent bootstrap probability values (≥30 %) for 1,000 replicates.

**Fig. S6b.**
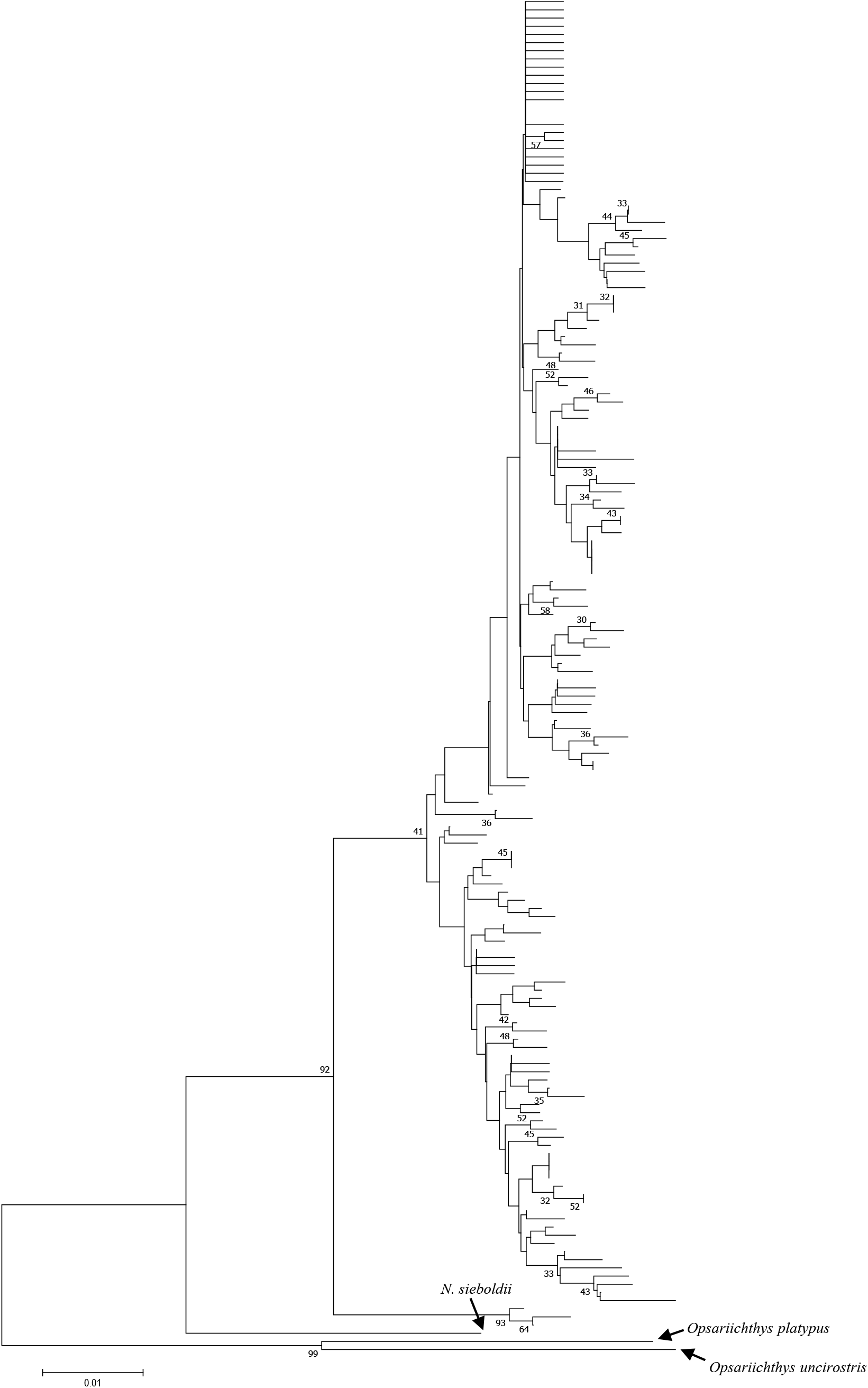
*N. temminckii*: NJ tree based on detected haplotypes after data screening step 1. Numbers at internodes represent bootstrap probability values (≥30 %) for 1,000 replicates.

**Fig. S6c.**
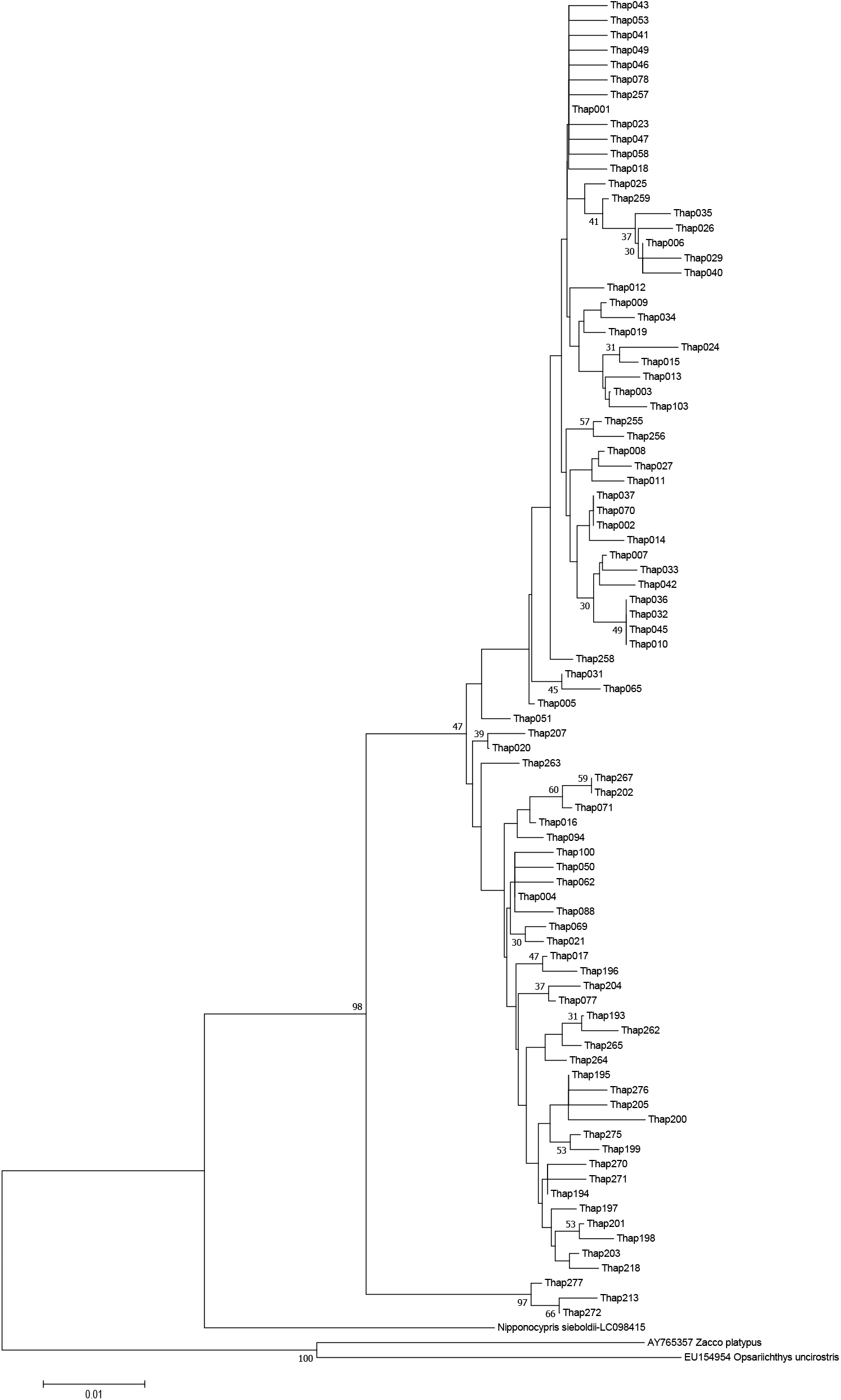
*N. temminckii*: NJ tree based on detected haplotypes after data screening step 2. Numbers at internodes represent bootstrap probability values (≥30 %) for 1,000 replicates.

**Fig. S6d.**
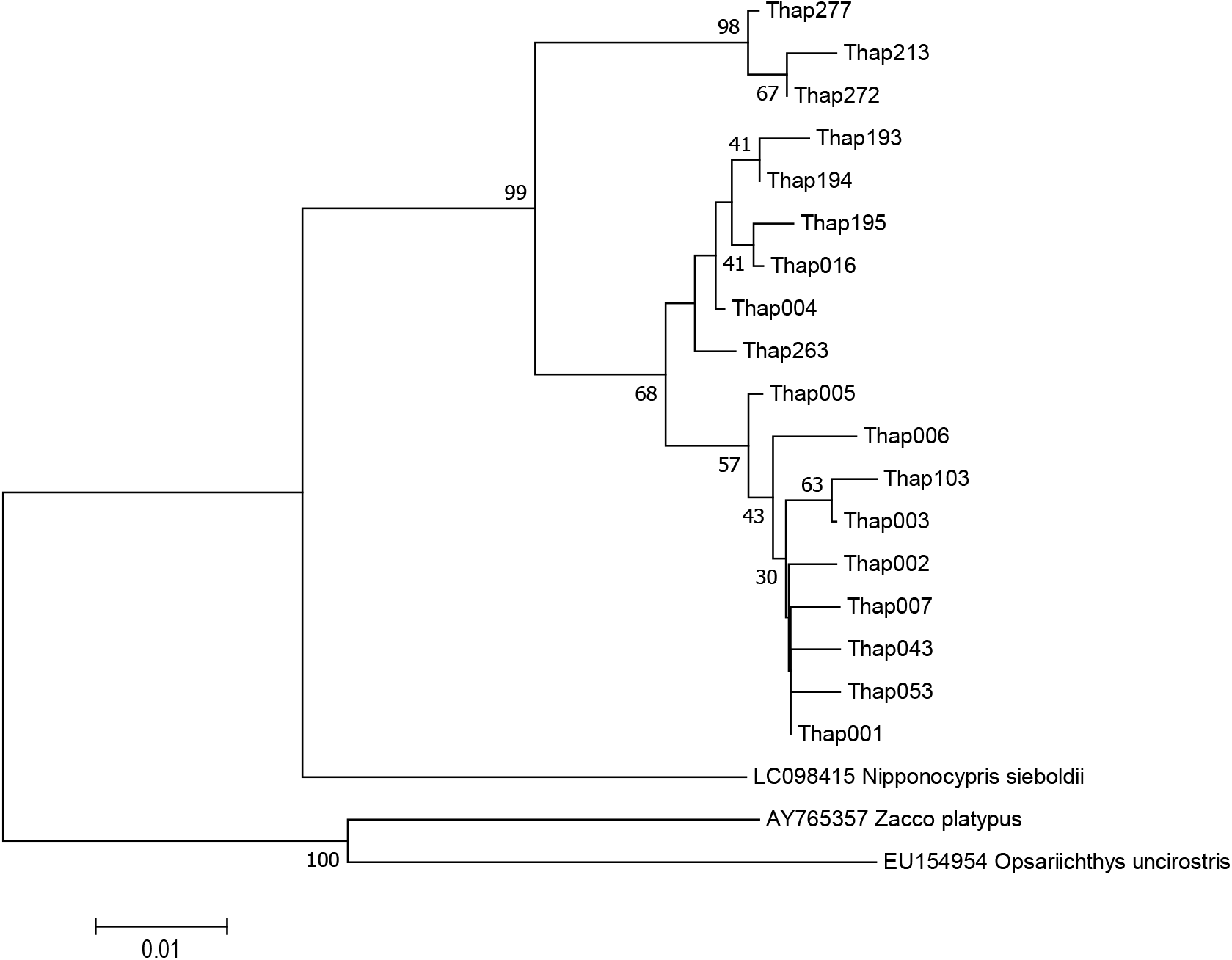
*N. temminckii*: NJ tree based on detected haplotypes after data screening step 3_1/3 (<max%/3 → 0%). Numbers at internodes represent bootstrap probability values (≥ 30 %) for 1,000 replicates.

**Fig. S7a.**
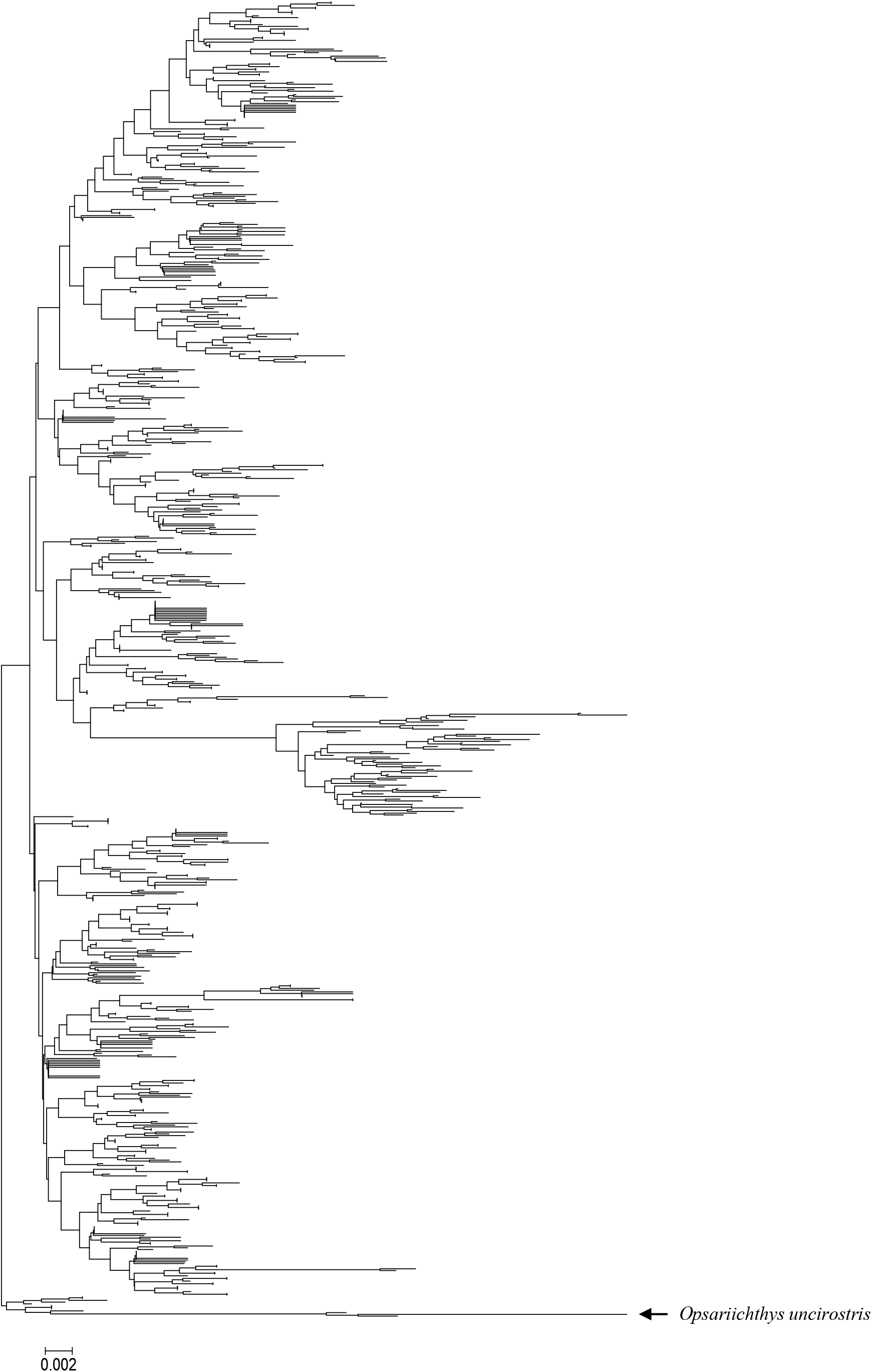
*Zacco platypus*: NJ tree based on detected haplotypes (step 0, non-data screening).

**Fig. S7b.**
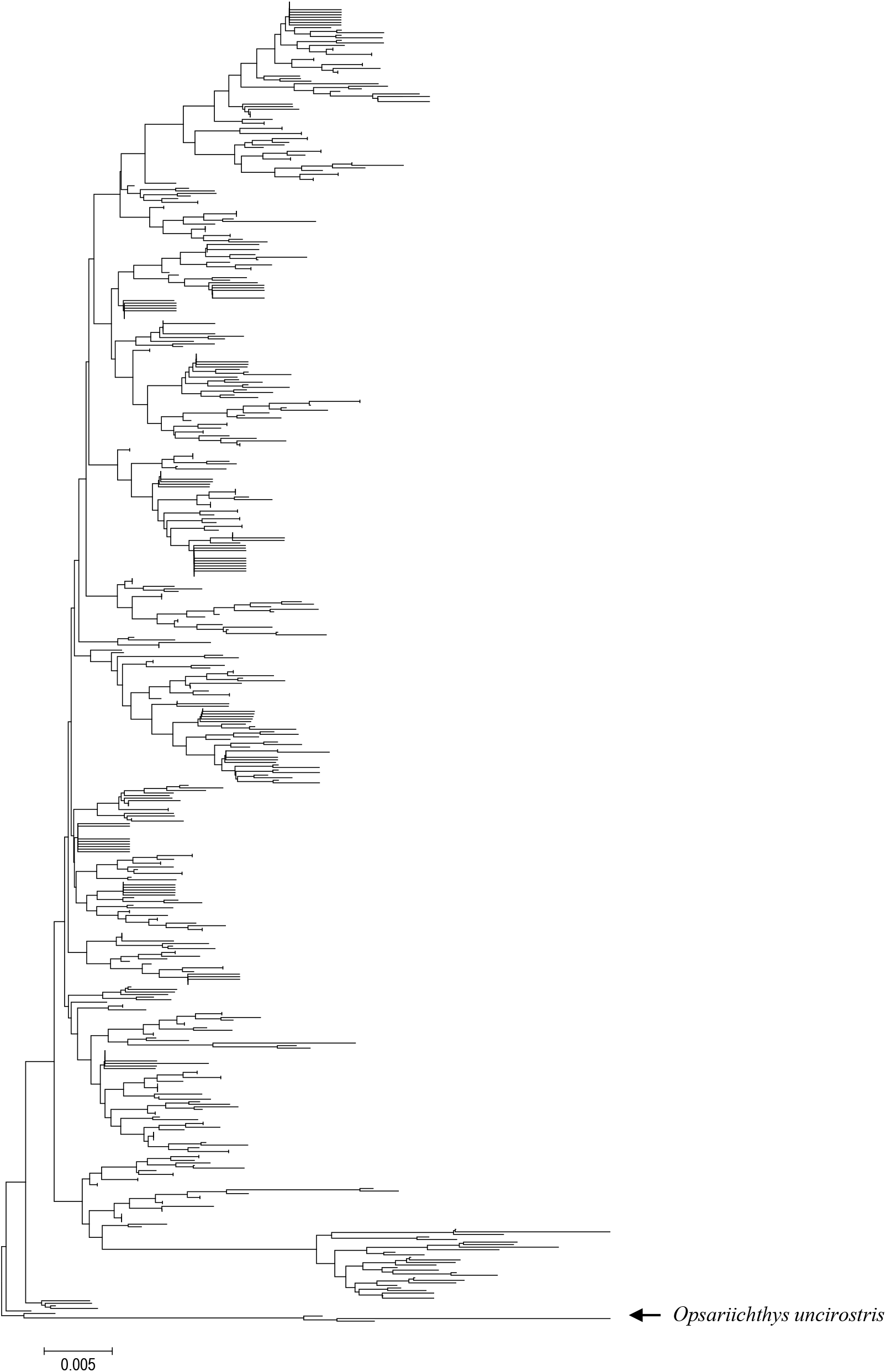
*Zaccoplatypus*:NJ tree based on detected haplotypes after data screening step 1.

**Fig. S7c.**
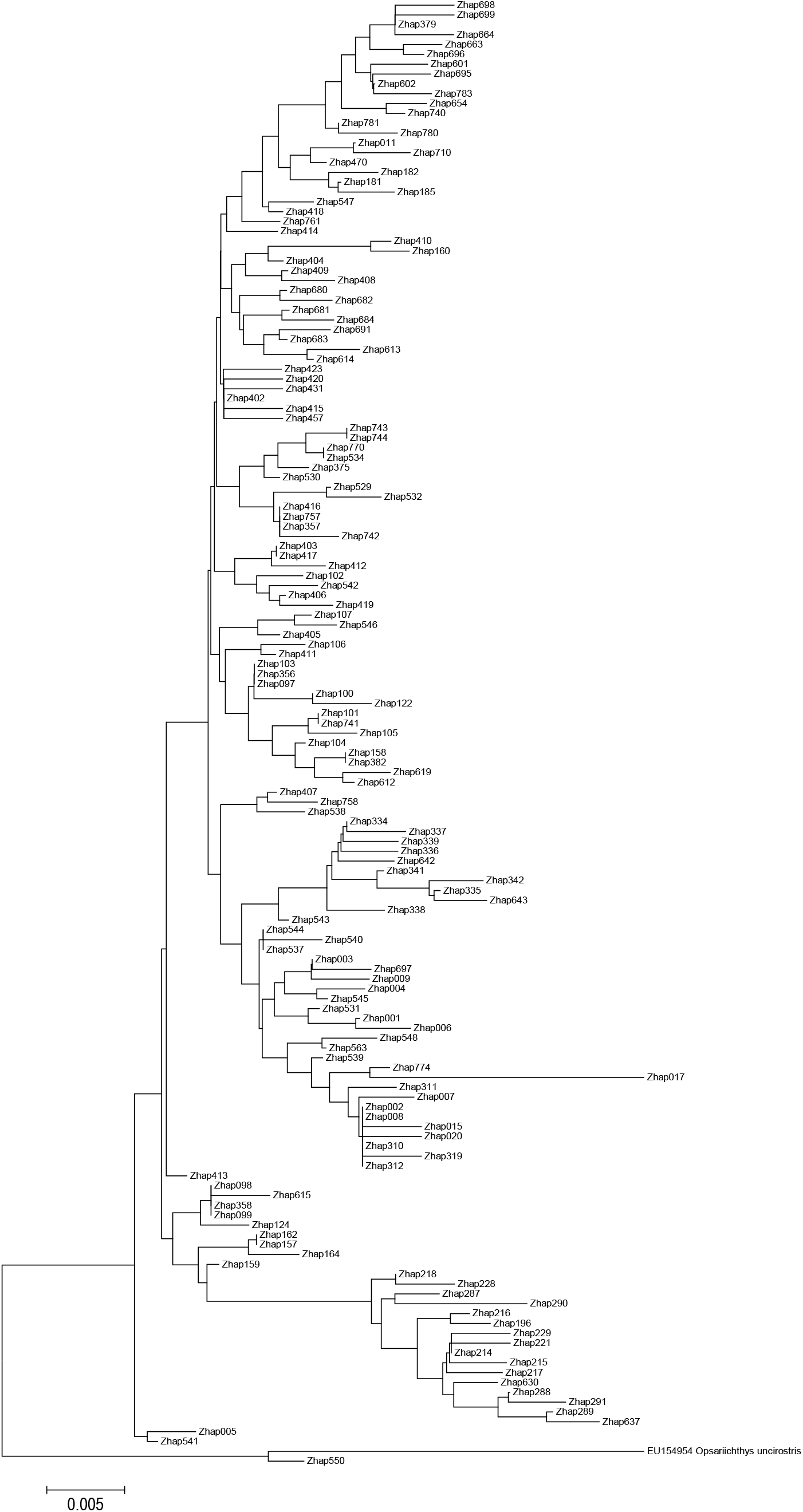
*Zaccoplatypus*: NJ tree based on detected haplotypes after data screening step 2.

**Fig. S7d.**
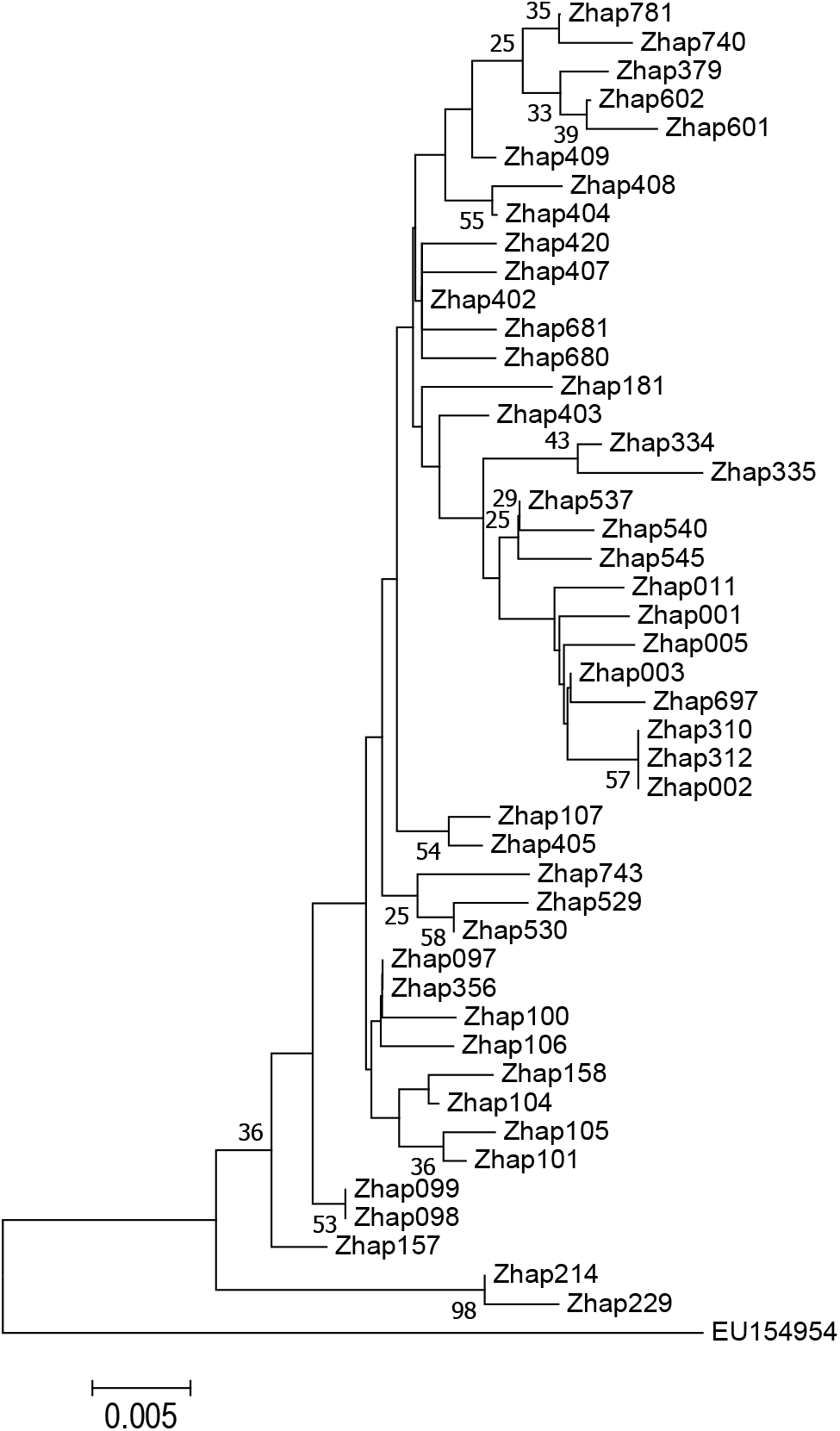
*Zacco platypus*: NJ tree based on detected haplotypes after data screening step 3_1/3 (<max%/3 → 0%). Numbers at internodes represent bootstrap probability values (≥ 30 %) for 1,000 replicates.

**Fig. S8.**
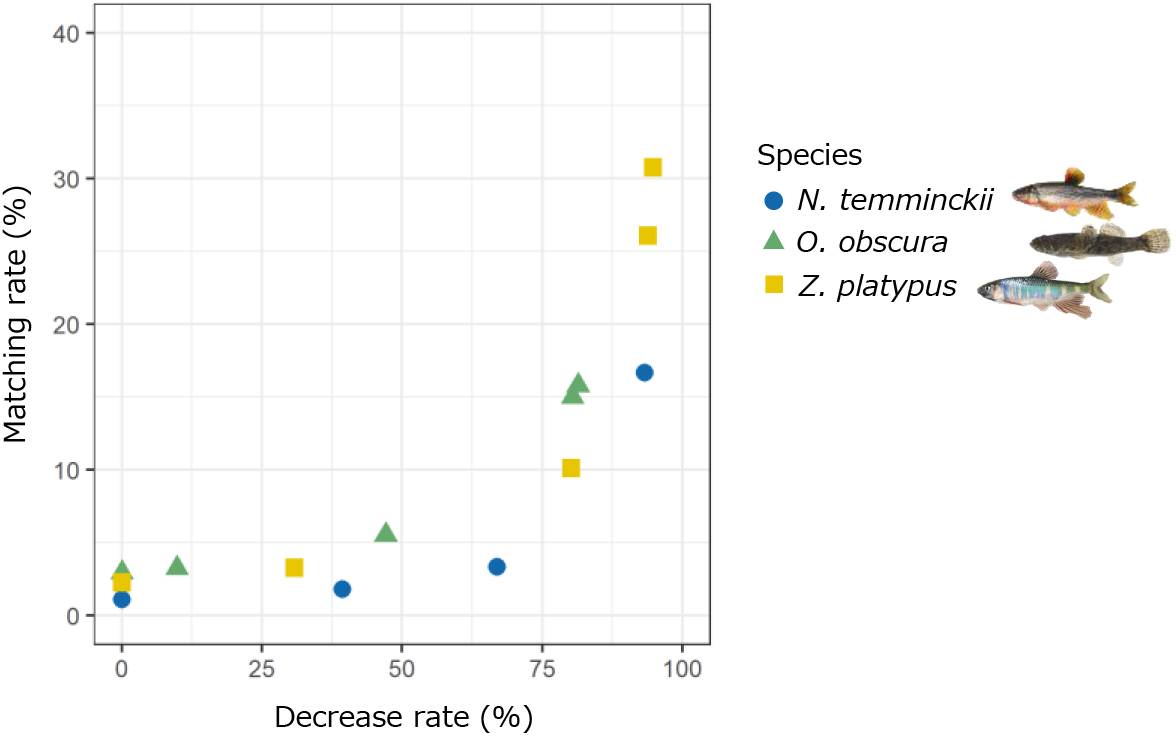
Relationship between the decrease rate of number of haplotypes from step 0 in data screening and matching rate with reference haplotypes (*p* < 0.001, GLMM).

**Fig. S9.**
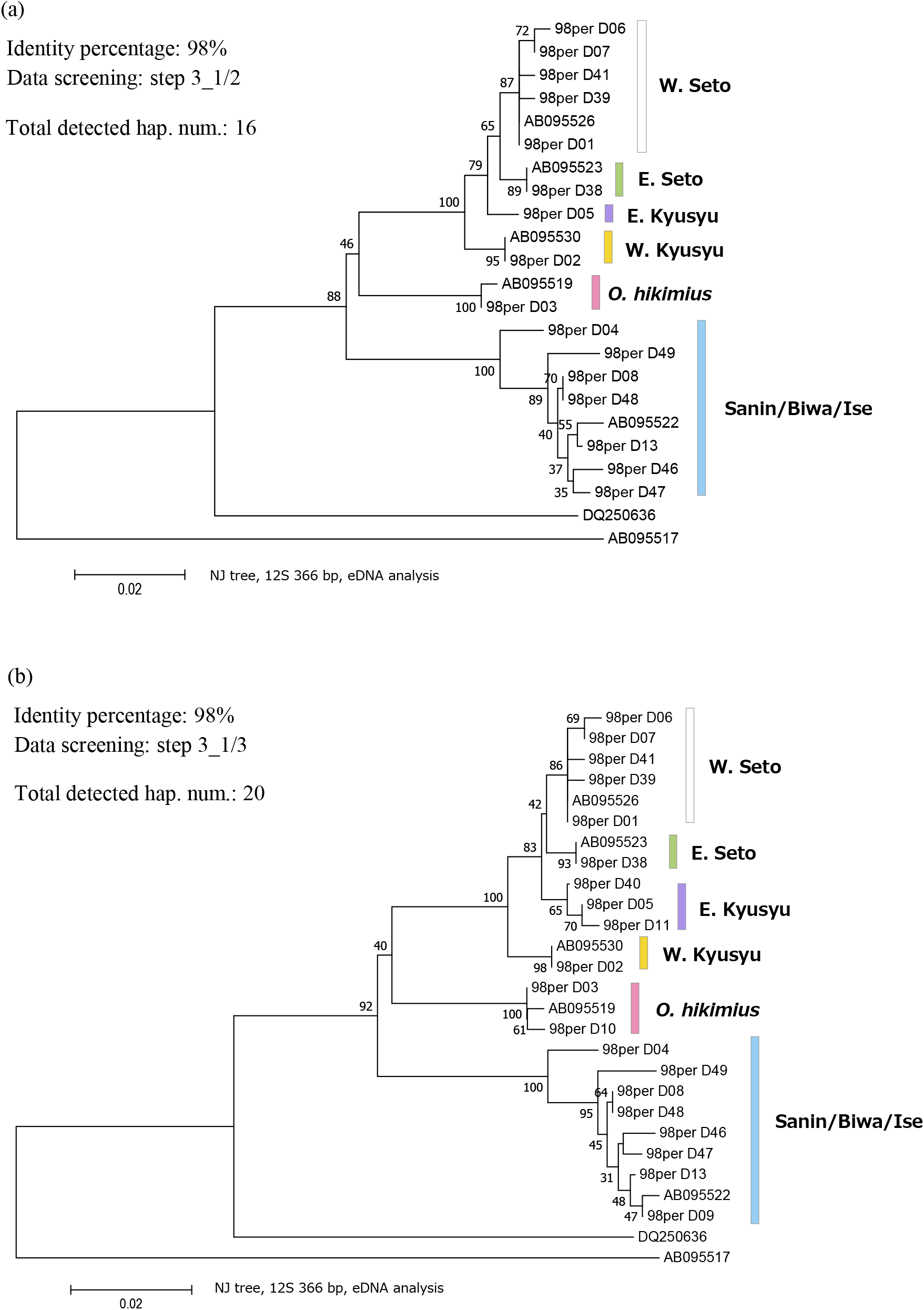
*Odontobutis obscura* and *O. hikimius*: Re-analysed with a 98% identity percentage in local BLASTN search. NJ tree based on the obtained sequence in (a) data screening step 3_1/2 and (b) 3_1/3. Numbers at internodes of NJ tree represent bootstrap probability values (≥ 30 %) for 1,000 replicates. The colours of each group are common in the panels.

**Fig. S10.**
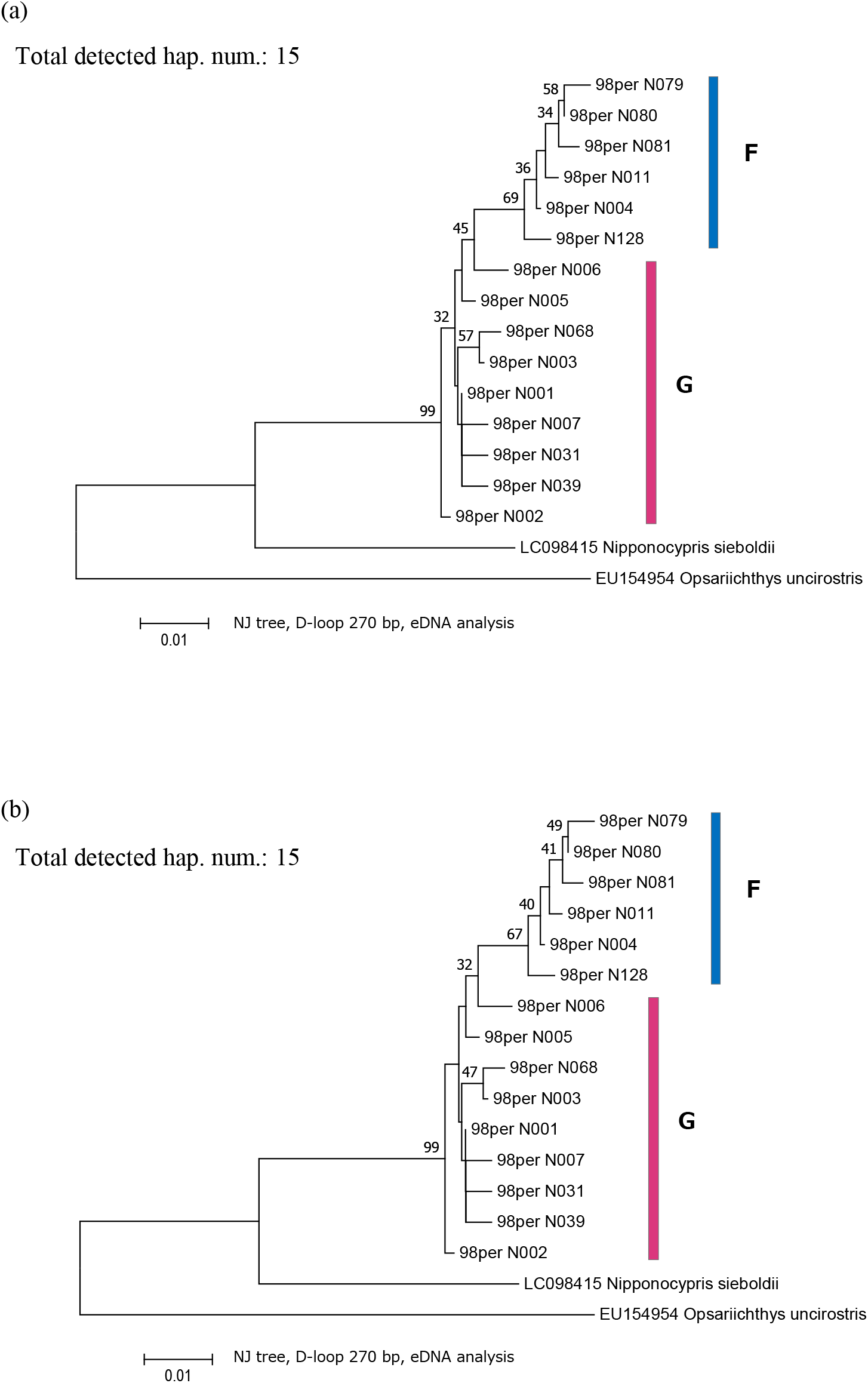
*Nipponocypris temminckii* : Re-analysed with a 98% identity percentage in local BLASTN search. NJ tree based on the obtained sequence in (a) data screening step 3_1/2 and (b) 3_1/3. Numbers at internodes of NJ tree represent bootstrap probability values (≥ 30 %) for 1,000 replicates. The colours of each group are common in the panels.

**Fig. S11.**
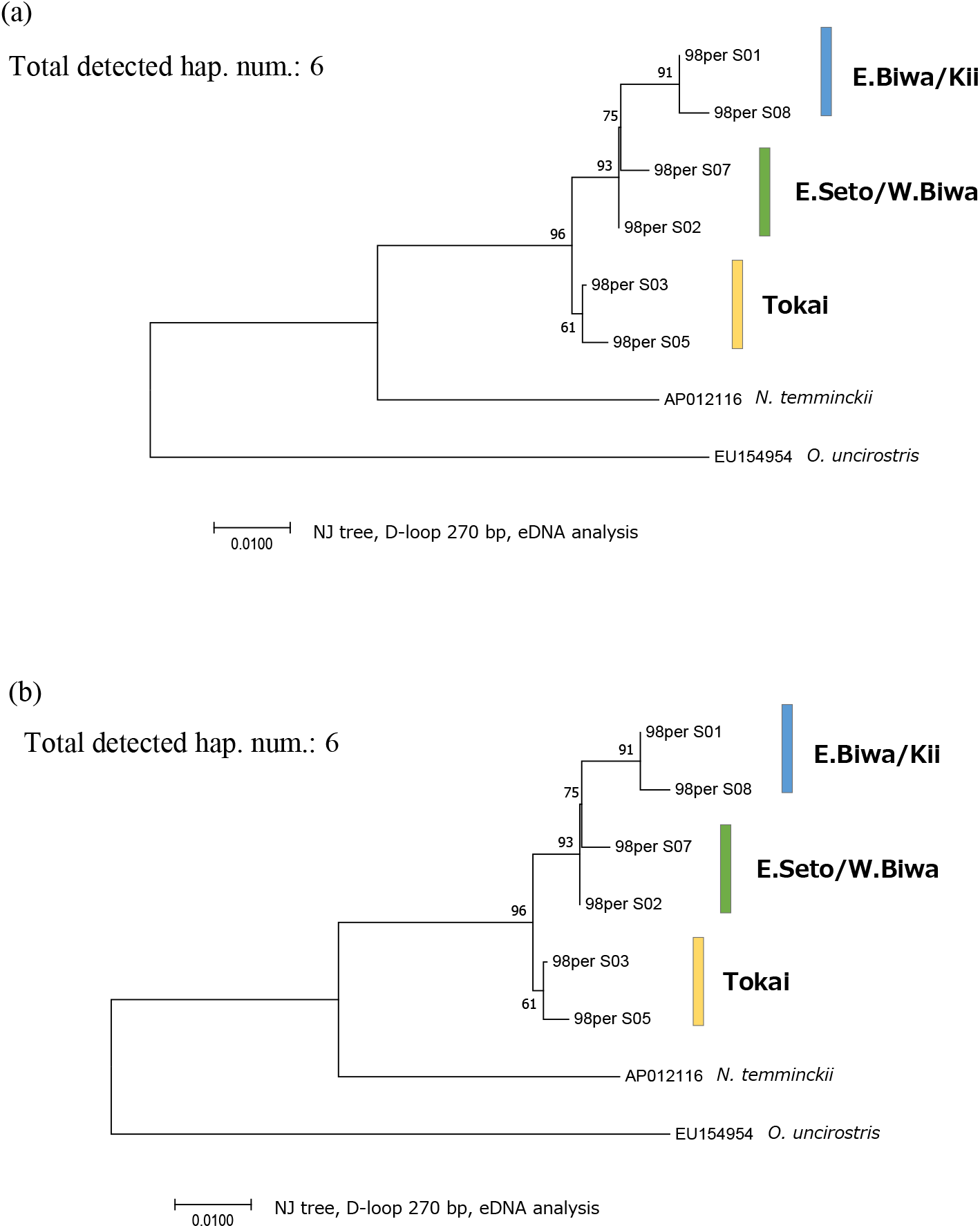
*Nipponocypris sieboldii* : Re-analysed with a 98% identity percentage in local BLASTN. NJ tree based on the obtained sequence in (a) data screening step 3_1/2 and (b) 3_1/3. Numbers at internodes of NJ tree represent bootstrap probability values (≥ 30 %) for 1,000 replicates. The colours of each group are common in the panels.

**Fig. S12.**
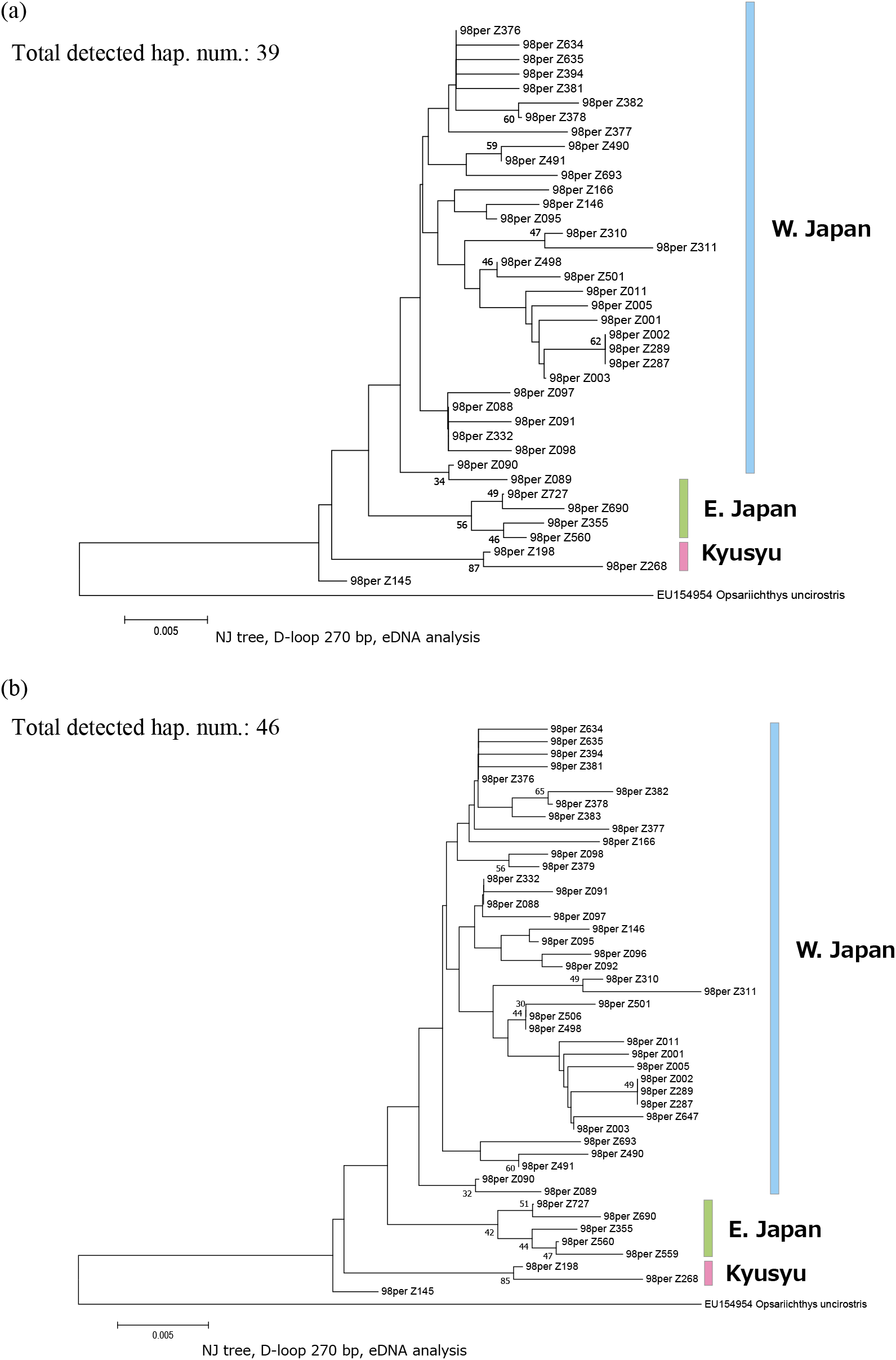
*Zacco platypus*: Re-analysed with a 98% identity percentage in local BLASTN. NJ tree based on the obtained sequence in (a) data screening step 3_1/2 and (b) 3_1/3. Numbers at internodes of NJ tree represent bootstrap probability values (≥ 30 %) for 1,000 replicates. The colours of each group are common in the panels.

## References

Andres, K.J., Sethi, S.A., Lodge, D.M., Andrés, J. (2021) Nuclear eDNA estimates population allele frequencies and abundance in experimental mesocosms and field samples. Molecular Ecology 30, 685–697. https://doi.org/10.1111/mec.15765

Aoyama, J., Watanabe, S., Ishikawa, S., Nishida, M., Tsukamoto, K (2000) Are morphological characters distinctive enough to discriminate between two species of freshwater eels,anguilla celebesensis andA. interioris? Ichthyological Research 47, 157–161. https://doi.org/10.1007/BF02684236

Avise, J.C. (2000) Phylogeography: The History and Formation of Species. Harvard University Press, Cambridge.

Avise, J.C., Arnold, J., Ball, R.M., Bermingham, E., Lamb, T., Neigel, J.E., Reeb, C.A., Saunders, N.C. (1987) Intraspecific phylogeography: the mitochondrial DNA bridge between population genetics and systematics. Annual review of ecology and systematics 18, 489–522.

Baker, C.S., Steel, D., Nieukirk, S., Klinck, H. (2018) Environmental DNA (eDNA) from the wake of the whales: droplet digital PCR for detection and species identification. Frontiers in Marine Science 5.

Ballard, J.W.O., Whitlock, M.C. (2004) The incomplete natural history of mitochondria. Molecular Ecology 13, 729–744. https://doi.org/10.1046/j.1365-294X.2003.02063.x

Bates, D., Maechler, M., Bolker, B., Walker, S., Christensen, R.H.B., Singmann, H., Dai, B., Scheipl, F., Grothendieck, G., Green, P., Fox, J., Bauer, A., Krivitsky, P.N. (2022) Linear Mixed-Effects Models using “Eigen” and S4. – R package ver.1.1-29.

Becker, R.A., Wilks, A.R., Brownrigg, R., Minka, T.P., Deckmyn, A. (2021) maps: Draw geographical maps. ver. 3.4.0.

Bermingham, E., Moritz, C. (1998) Comparative phylogeography: concepts and applications. Molecular Ecology 7, 367–369. https://doi.org/10.1046/j.1365-294x.1998.00424.x

Brownrigg, R., 2018. Extra Map Databases. Version 2.3.0 [R package].

Bylemans, J., Furlan, E.M., Hardy, C.M., McGuffie, P., Lintermans, M., Gleeson, D.M. (2017) An environmental DNA-based method for monitoring spawning activity: A case study, using the endangered Macquarie perch (Macquaria australasica). Methods in Ecology and Evolution 8, 646–655.

Callahan, B.J., McMurdie, P.J., Rosen, M.J., Han, A.W., Johnson, A.J.A., Holmes, S.P. (2016) DADA2: High-resolution sample inference from Illumina amplicon data. Nat Methods 13, 581–583. https://doi.org/10.1038/nmeth.3869

Camacho, C., Coulouris, G., Avagyan, V., Ma, N., Papadopoulos, J., Bealer, K., Madden, T.L. (2009) BLAST+: architecture and applications. BMC Bioinformatics 10, 421. https://doi.org/10.1186/1471-2105-10-421

Corush, J.B., Pierson, T.W., Shiao, J.-C., Katayama, Y., Zhang, J., Fitzpatrick, B.M. (2022) Amphibious mudskipper populations are genetically connected along coastlines, but differentiated across water. Journal of Biogeography 49, 767–779. https://doi.org/10.1111/jbi.14345

Cucherousset, J., Olden, J.D. (2011) Ecological impacts of nonnative freshwater fishes. Fisheries 36, 215–230. https://doi.org/10.1080/03632415.2011.574578

Deiner, K., Renshaw, M.A., Li, Y., Olds, B.P., Lodge, D.M., Pfrender, M.E. (2017) Long-range PCR allows sequencing of mitochondrial genomes from environmental DNA. Methods in Ecology and Evolution 8, 1888–1898. https://doi.org/10.1111/2041-210X.12836

Dugal, L., Thomas, L., Jensen, M.R., Sigsgaard, E.E., Simpson, T., Jarman, S., Thomsen, P.F., Meekan, M. (2022) Individual haplotyping of whale sharks from seawater environmental DNA. Molecular Ecology Resources 22, 56–65. https://doi.org/10.1111/1755-0998.13451

Edgar, R.C. (2016) UNOISE2: improved error-correction for Illumina 16S and ITS amplicon sequencing. bioRxiv. https://doi.org/10.1101/081257

Gerritsen, H. (2018) Data Visualisation on Maps. ver. 1.5.1.

Gozlan, R.E., Britton, J.R., Cowx, I., Copp, G.H. (2010) Current knowledge on non-native freshwater fish introductions. Journal of Fish Biology 76, 751–786. https://doi.org/10.1111/j.1095-8649.2010.02566.x

Hamady, M., Walker, J.J., Harris, J.K., Gold, N.J., Knight, R. (2008) Error-correcting barcoded primers for pyrosequencing hundreds of samples in multiplex. Nat Methods 5, 235–237. https://doi.org/10.1038/nmeth.1184

Holman, L.E., Parker-Nance, S., de Bruyn, M., Creer, S., Carvalho, G., Rius, M. (2022) Managing human-mediated range shifts: understanding spatial, temporal and genetic variation in marine non-native species. Philosophical Transactions of the Royal Society B: Biological Sciences 377, 20210025. https://doi.org/10.1098/rstb.2021.0025

Hosoya, K. (2019) Sankei Handy Illustrated Book 15: Freshwater fish of Japan, enlarged and revised edition. ed. Yama-kei Publishers co.,Ltd.

Iwata, A., Sakai, H. (2002) Odontobutis hikimius: A new freshwater goby from Japan, with a key to species of the Genus. cope 2002, 104–110. https://doi.org/10.1643/0045-8511(2002)002[0104:OHNSAN]2.0.CO;2

Jensen, M.R., Sigsgaard, E.E., Liu, S., Manica, A., Bach, S.S., Hansen, M.M., Møller, P.R., Thomsen, P.F. (2021) Genome-scale target capture of mitochondrial and nuclear environmental DNA from water samples. Molecular Ecology Resources 21, 690–702. https://doi.org/10.1111/1755-0998.13293

Jo, T., Murakami, H., Masuda, R., Sakata, M.K., Yamamoto, S., Minamoto, T. (2017) Rapid degradation of longer DNA fragments enables the improved estimation of distribution and biomass using environmental DNA. Molecular Ecology Resources 17, e25–e33. https://doi.org/10.1111/1755-0998.12685

Jukes, T.H., Cantor, C.R. (1969) Evolution of Protein Molecules. In: Munro, H.N., Ed., Mammalian Protein Metabolism. Academic Press, New York.

Kakehashi, R., Ito, S., Yasui, K., Kambayashi, Ch., Kanao, Sh., Kurabayashi, A. (2022) Amplification and sequencing of the complete mtDNA of the endangered bitterling, Acheilognathus longipinnis (Cyprinidae), using environmental DNA from aquarium water. J. Ichthyol. 62, 280–288. https://doi.org/10.1134/S0032945222020072

Kitanishi, S., Hayakawa, A., Takamura, K., Nakajima, J., Kawaguchi, Y., Onikura, N., Mukai, T. (2016) Phylogeography of Opsariichthys platypus in Japan based on mitochondrial DNA sequences. Ichthyol Res 63, 506–518. https://doi.org/10.1007/s10228-016-0522-y

Marshall, N.T., Stepien, C.A. (2019) Invasion genetics from eDNA and thousands of larvae: A targeted metabarcoding assay that distinguishes species and population variation of zebra and quagga mussels. Ecology and Evolution 9, 3515–3538. https://doi.org/10.1002/ece3.4985

Miya, M., Sato, Y., Fukunaga, T., Sado, T., Poulsen, J.Y., Sato, K., Minamoto, T., Yamamoto, S., Yamanaka, H., Araki, H. (2015) MiFish, a set of universal PCR primers for metabarcoding environmental DNA from fishes: detection of more than 230 subtropical marine species. Royal Society open science 2, 150088.

Miyake, T., Nakajima, J., Onikura, N., Ikemoto, S., Iguchi, K., Komaru, A., Kawamura, K. (2011) The genetic status of two subspecies of Rhodeus atremius, an endangered bitterling in Japan. Conserv Genet 12, 383–400. https://doi.org/10.1007/s10592-010-0146-0

Miyake, T., Nakajima, J., Umemura, K., Onikura, N., Ueda, T., Smith, C., Kawamura, K. (2021) Genetic diversification of the Kanehira bitterling Acheilognathus rhombeus inferred from mitochondrial DNA, with comments on the phylogenetic relationship with its sister species Acheilognathus barbatulus. Journal of Fish Biology 99, 1677–1695. https://doi.org/10.1111/jfb.14876

Miyazaki, J.-I., Dobashi, M., Tamura, T., Beppu, S., Sakai, T., Mihara, M., Hosoya, K. (2011) Parallel evolution in eight-barbel loaches of the genus Lefua (Balitoridae, Cypriniformes) revealed by mitochondrial and nuclear DNA phylogenies. Molecular Phylogenetics and Evolution 60, 416–427. https://doi.org/10.1016/j.ympev.2011.05.005

Mizuguchi, K. (1990) Dispersal of the oikawa, Zacco platypus (temminck et schlegel), in Japan. Rep Tokyo Univ Fish 25, 149–169.

Moritz, C. (2002) Strategies to protect biological diversity and the evolutionary processes that sustain it. Systematic Biology 51, 238–254. https://doi.org/10.1080/10635150252899752

Mukai, T., Nishida, M. (2003) Mitochondrial DNA phylogeny of Japanese freshwater goby, Odontobutis obscura, and an evidence for artificial transplantation to Kanto District. Japan. J. Ichthyol. 50, 71–76.

Mukai, T., Onikura, N., Yodo, T., Senou, H. (2013) Domestic alien fishes: Hidden threats to biodiversity, Edited by Nature Conservation Committee of Ichtyological Society of Japan. ed. Tokai University Press.

Nakagawa, H., Seki, S., Ishikawa, T., Watanabe, K. (2016) Genetic population structure of the Japanese torrent catfish Liobagrus reinii (Amblycipitidae) inferred from mitochondrial cytochrome b variations. Ichthyol Res 63, 333–346. https://doi.org/10.1007/s10228-015-0503-6

Nguyen, T.V., Tilker, A., Nguyen, A., Hörig, L., Axtner, J., Schmidt, A., Le, M., Nguyen, A.H.Q., Rawson, B.M., Wilting, A., Fickel, J. (2021) Using terrestrial leeches to assess the genetic diversity of an elusive species: The Annamite striped rabbit Nesolagus timminsi. Environmental DNA 3, 780–791. https://doi.org/10.1002/edn3.182

Palumbi, S., Martin, A., Romano, S., McMillian, W., Stice, L., Grabowski, G. (1991) The simple fool’s guide to PCR. Univ Hawaii, Honolulu.

Parsons, K.M., Everett, M., Dahlheim, M., Park, L. (2018) Water, water everywhere: environmental DNA can unlock population structure in elusive marine species. Royal Society Open Science 5, 180537. https://doi.org/10.1098/rsos.180537

R Core Team. R, 2021. A Language and Environment for Statistical Computing.

Rees, H.C., Maddison, B.C., Middleditch, D.J., Patmore, J.R.M., Gough, K.C. (2014) The detection of aquatic animal species using environmental DNA – a review of eDNA as a survey tool in ecology. Journal of Applied Ecology 51, 1450–1459. https://doi.org/10.1111/1365-2664.12306

Ruzzante, D.E., Walde, S.J., Gosse, J.C., Cussac, V.E., Habit, E., Zemlak, T.S., Adams, E.D.M. (2008) Climate control on ancestral population dynamics: insight from Patagonian fish phylogeography. Molecular Ecology 17, 2234–2244. https://doi.org/10.1111/j.1365-294X.2008.03738.x

Saitou, N., Nei, M. (1987). The neighbor-joining method: a new method for reconstructing phylogenetic trees. Molecular Biology and Evolution 4, 406–425. https://doi.org/10.1093/oxfordjournals.molbev.a040454

Sakai, H., Yamamoto, C., Iwata, A. (1998) Genetic divergence, variation and zoogeography of a freshwater goby,Odontobutis obscura. Ichthyol Res 45, 363–376. https://doi.org/10.1007/BF02725189

Scoble, J., Lowe, A.J. (2010) A case for incorporating phylogeography and landscape genetics into species distribution modelling approaches to improve climate adaptation and conservation planning. Diversity and Distributions 16, 343–353. https://doi.org/10.1111/j.1472-4642.2010.00658.x

Sersics, A.N., Cosacov, A., Cocucci, A.C., Johnson, L.A., Pozner, R., Avila, L.J., Sites, J.W., Jr., Morando, M. (2011) Emerging phylogeographical patterns of plants and terrestrial vertebrates from Patagonia. Biological Journal of the Linnean Society 103, 475–494. https://doi.org/10.1111/j.1095-8312.2011.01656.x

Shum, P., Palumbi, S.R. (2021) Testing small-scale ecological gradients and intraspecific differentiation for hundreds of kelp forest species using haplotypes from metabarcoding. Molecular Ecology 30, 3355–3373. https://doi.org/10.1111/mec.15851

Sigsgaard, E.E., Jensen, M.R., Winkelmann, I.E., Møller, P.R., Hansen, M.M., Thomsen, P.F. (2020) Population-level inferences from environmental DNA—Current status and future perspectives. Evol Appl 13, 245–262. https://doi.org/10.1111/eva.12882

Sigsgaard, E.E., Nielsen, I.B., Bach, S.S., Lorenzen, E.D., Robinson, D.P., Knudsen, S.W., Pedersen, M.W., Jaidah, M.A., Orlando, L., Willerslev, E., Møller, P.R., Thomsen, P.F. (2016) Population characteristics of a large whale shark aggregation inferred from seawater environmental DNA. Nat Ecol Evol 1, 1–5. https://doi.org/10.1038/s41559-016-0004

Soltis, D.E., Morris, A.B., McLACHLAN, J.S., Manos, P.S., Soltis, P.S. (2006) Comparative phylogeography of unglaciated eastern North America. Molecular Ecology 15, 4261–4293. https://doi.org/10.1111/j.1365-294X.2006.03061.x

Stat, M., Huggett, M.J., Bernasconi, R., DiBattista, J.D., Berry, T.E., Newman, S.J., Harvey, E.S., Bunce, M. (2017) Ecosystem biomonitoring with eDNA: metabarcoding across the tree of life in a tropical marine environment. Sci Rep 7, 12240. https://doi.org/10.1038/s41598-017-12501-5

Székely, D., Corfixen, N.L., Mørch, L.L., Knudsen, S.W., McCarthy, M.L., Teilmann, J., Heide-Jørgensen, M.P., Olsen, M.T. (2021) Environmental DNA captures the genetic diversity of bowhead whales (Balaena mysticetus) in West Greenland. Environmental DNA 3, 248–260. https://doi.org/10.1002/edn3.176

Taberlet, P., Coissac, E., Hajibabaei, M., Rieseberg, L.H. (2012) Environmental DNA. Molecular Ecology 21, 1789–1793. https://doi.org/10.1111/j.1365-294X.2012.05542.x

Takamura, K., Nakahara, M. (2015) Intraspecific invasion occurring in geographically isolated populations of the Japanese cyprinid fish Zacco platypus. Limnology 16, 161–170. https://doi.org/10.1007/s10201-015-0450-y

Takehana, Y., Nagai, N., Matsuda, M., Tsuchiya, K., Sakaizumi, M. (2003) Geographic variation and diversity of the Cytochrome b gene in Japanese wild populations of Medaka, Oryzias latipes. jzoo 20, 1279–1291. https://doi.org/10.2108/zsj.20.1279

Taniguchi, S., Bertl, J., Futschik, A., Kishino, H., Okazaki, T. (2021) Waves out of the Korean Peninsula and inter- and intra-species replacements in freshwater fishes in Japan. Genes 12, 303. https://doi.org/10.3390/genes12020303

Teske, P.R., Golla, T.R., Sandoval-Castillo, J., Emami-Khoyi, A., van der Lingen, C.D., von der Heyden, S., Chiazzari, B., Jansen van Vuuren, B., Beheregaray, L.B. (2018) Mitochondrial DNA is unsuitable to test for isolation by distance. Sci Rep 8, 8448. https://doi.org/10.1038/s41598-018-25138-9

Thomsen, P.F., Willerslev, E. (2015) Environmental DNA–An emerging tool in conservation for monitoring past and present biodiversity. Biological conservation 183, 4–18.

Tominaga, K., Nagata, N., Kitamura, J., Watanabe, K., Sota, T. (2020) Phylogeography of the bitterling Tanakia lanceolata (Teleostei: Cyprinidae) in Japan inferred from mitochondrial cytochrome b gene sequences. Ichthyol Res 67, 105–116. https://doi.org/10.1007/s10228-019-00715-8

Tominaga, K., Nakajima, J., Watanabe, K. (2016) Cryptic divergence and phylogeography of the pike gudgeon Pseudogobio esocinus (Teleostei: Cyprinidae): a comprehensive case of freshwater phylogeography in Japan. Ichthyol Res 63, 79–93. https://doi.org/10.1007/s10228-015-0478-3

Tsuji, S., Inui, R., Nakao, R., Miyazono, S., Saito, M., Kono, T., Akamatsu, Y. (2022a) Quantitative environmental DNA metabarcoding reflects quantitative capture data of fish community obtained by electrical shocker. bioRxiv, https://doi.org/10.1101/2022.04.27.489619

Tsuji, S., Maruyama, A., Miya, M., Ushio, M., Sato, H., Minamoto, T., Yamanaka, H. (2020a) Environmental DNA analysis shows high potential as a tool for estimating intraspecific genetic diversity in a wild fish population. Molecular Ecology Resources 20, 1248–1258.

Tsuji, S., Miya, M., Ushio, M., Sato, H., Minamoto, T., Yamanaka, H. (2020b) Evaluating intraspecific genetic diversity using environmental DNA and denoising approach: A case study using tank water. Environmental DNA 2, 42–52. https://doi.org/10.1002/edn3.44

Tsuji, S., Murakami, H., Masuda, R. (2022b) Analysis of the persistence and particle size distributional shift of sperm-derived environmental DNA to monitor Jack Mackerel spawning activity. https://doi.org/10.1101/2022.03.09.483695

Tsuji, S., Shibata, N. (2021) Identifying spawning events in fish by observing a spike in environmental DNA concentration after spawning. Environmental DNA 3, 190–199.

Tsuji, S., Shibata, N., Sawada, H., Ushio, M. (2020c) Quantitative evaluation of intraspecific genetic diversity in a natural fish population using environmental DNA analysis. Molecular Ecology Resources 20, 1323–1332.

Turon, X., Antich, A., Palacín, C., Præbel, K., Wangensteen, O.S. (2020) From metabarcoding to metaphylogeography: separating the wheat from the chaff. Ecological Applications 30, e02036. https://doi.org/10.1002/eap.2036

Ushio, M., Murakami, H., Masuda, R., Sado, T., Miya, M., Sakurai, S., Yamanaka, H., Minamoto, T., Kondoh, M. (2018) Quantitative monitoring of multispecies fish environmental DNA using high-throughput sequencing. Metabarcoding and Metagenomics 2, e23297. https://doi.org/10.3897/mbmg.2.23297

Watanabe, K., Mori, S., Tanaka, T., Kanagawa, N., Itai, T., Kitamura, J., Suzuki, N., Tominaga, K., Kakioka, R., Tabata, R., Abe, T., Tashiro, Y., Hashimoto, Y., Nakajima, J., Onikura, N. (2014) Genetic population structure of Hemigrammocypris rasborella (Cyprinidae) inferred from mtDNA sequences. Ichthyol Res 61, 352–360. https://doi.org/10.1007/s10228-014-0406-y

Watanabe, K., Takahashi, H., Kitamura, A., Yokoyama, R., Kitagawa, T., Takeshima, H., Sato, S., Yamamoto, S., Yusuke, T., Mikai, T., Ohara, K., Iguchi, K. (2006) Biogeographical history of Japanese freshwater fishes: Phylogeographic approaches and perspectives. Japanese journal of Ichthyology 53, 1–38. https://doi.org/10.11369/jji1950.53.1

Weitemier, K., Penaluna, B.E., Hauck, L.L., Longway, L.J., Garcia, T., Cronn, R. (2021) Estimating the genetic diversity of Pacific salmon and trout using multigene eDNA metabarcoding. Molecular Ecology 30, 4970–4990. https://doi.org/10.1111/mec.15811

Yamamoto, S., Masuda, R., Sato, Y., Sado, T., Araki, H., Kondoh, M., Minamoto, T., Miya, M. (2017) Environmental DNA metabarcoding reveals local fish communities in a species-rich coastal sea. Sci Rep 7, 40368. https://doi.org/10.1038/srep40368

Yamanaka, H., Minamoto, T., Matsuura, J., Sakurai, S., Tsuji, S., Motozawa, H., Hongo, M., Sogo, Y., Kakimi, N., Teramura, I., Sugita, M., Baba, M., Kondo, A. (2017) A simple method for preserving environmental DNA in water samples at ambient temperature by addition of cationic surfactant. Limnology 18, 233–241. https://doi.org/10.1007/s10201-016-0508-5

Yoshitake, K., Yoshinaga, T., Tanaka, C., Mizusawa, N., Reza, Md.S., Tsujimoto, A., Kobayashi, T., Watabe, S. (2019) HaCeD-Seq: a Novel Method for Reliable and Easy Estimation About the Fish Population Using Haplotype Count from eDNA. Mar Biotechnol 21, 813–820. https://doi.org/10.1007/s10126-019-09926-6

Zizka, V.M.A., Koschorreck, J., Khan, C.C., Astrin, J.J. (2022) Long-term archival of environmental samples empowers biodiversity monitoring and ecological research. Environmental Sciences Europe 34, 40. https://doi.org/10.1186/s12302-022-00618-y

