## Appendix for "Environmental DNA phylogeography: successful reconstruction of phylogeographic patterns of multiple fish species from a cup of water"

### Abstract

Phylogeography is an integrative field of science linking micro- and macro-evolutionary processes, contributing to the inference of vicariance, dispersal, speciation, and other population-level processes. Phylogeographic surveys usually require considerable effort and time to obtain numerous samples from many geographical sites covering the distribution range of target species; this associated high cost limits their application. Recently, environmental DNA (eDNA) analysis has been useful not only for detecting species but also for assessing genetic diversity; hence, there has been growing interest in its application to phylogeography. As the first step of eDNA-based phylogeography, we examined (1) data screening procedures suitable for phylogeography and (2) whether the results obtained from eDNA analysis accurately reflect known phylogeographic patterns. For these purposes, we performed quantitative eDNA metabarcoding using group-specific primer sets in five freshwater fish species belonging to two taxonomic groups from a total of 94 water samples collected from western Japan. As a result, three-step data screening based on the DNA copy number of each haplotype detected successfully eliminated suspected false positive haplotypes. Furthermore, eDNA analysis could almost perfectly reconstruct the phylogenetic and phylogeographic patterns obtained for all target species with the conventional method. Despite existing limitations and future challenges, eDNA-based phylogeography can significantly reduce survey time and effort and is applicable for simultaneous analysis of multiple species in single water samples. eDNA-based phylogeography has the potential to revolutionise phylogeography.

**Keywords:** comparative phylogeography, environmental DNA, freshwater fish

### Appendix: Further Methodological Details

#### *DNA extraction from the filter sample*

DNA extraction from filter samples was performed according to two different procedures: (a) Tsuji et al (2022a) for the 67 sites surveyed between 2017 and 2020; (b) Tsuji et al. (2022b) for the 27 sites surveyed in 2021.

(a) First, each filter sample was placed in the upper part of the Salivette tube (SARSTEDT AG & Co. KG, Nümbrecht, Germany). The 220 µL of extraction solution containing 200 µL Buffer AL and 20 µL of proteinase K in DNeasy Blood & Tissue Kit (Qiagen, Hilden, Germany) were added onto each filter and incubated at 56˚C for 30 min. After incubation, the Salivette tube was centrifuged at 5,000 × *g* for 1 min. 220 µL Tris–EDTA (TE) buffer (pH 8.0; Nippon Gene Co., Ltd., Tokyo, Japan) was added onto each filter and re-centrifuged at 5,000 × *g* for 3 min. After that, 200 µL of ethanol was added to the solution in the bottom part of the Salivette tube and mixed well by gently pipetting. The whole of the mixed solution was transferred to a DNeasy Mini spin column, and the DNA was purified according to the manufacturer's protocol. The DNA was finally eluted in 100 µL Buffer AE and stored at −20˚C.

(b) First, each filter sample was placed in the upper part of the spin column (EconoSpin, EP-31201; GeneDesign, Inc., Osaka, Japan) with its silica-gel membrane removed. Spin columns were centrifuged for 1 min at 6,000 × *g*, and any excess water contained in the filter was removed. A total of 420 µL of a solution, composed of 200 µL ultrapure water, 200 µL Buffer AL, and 20 µL proteinase K, was placed on each filter, and the spin columns were incubated for 45 min at 56 °C. After incubating, the spin columns were centrifuged for 1 min at 6,000 × *g*. 100 µL of Tris-EDTA buffer (pH 8.0) was placed on the filter and re-incubated for 1 min at room temperature. After centrifugation for 1 min at 6,000 × *g*, 600 µL of ethanol was added to collected liquid and mixed well by pipetting. The DNA mixture was transferred to a DNeasy mini spin column, and DNA was purified following the manufacturer's protocol. The DNA was finally eluted in 100 µL of Buffer AE.

#### *Sanger sequencing of tissue DNA*

Sanger sequencing of tissue DNA was performed on ten individuals of *Odontobutis obscura* caught in sts. 88 and 89 (Fukuchi River), where haplotypes belonging to an unreported lineage group (E. Kyushu group, see Results) were newly identified, and on *Nipponocypris sieboldii* collected from several sites in western Japan (Table S4).

(a) The mtDNA partial 12S sequence (777 bp) of *O. obscura* was amplified using the following primer pairs developed by ST: Odontobutis_sanger-F (5′-AGG GCC AGT AAA ACT CGT GC-3′) and Odontobutis_sanger-R (5′-GGG CGT CTT CTC GGT GTA AG-3′). The PCR was performed in a 20-µL total volume of reaction mixture containing 0.1 µL of TaKaRa Ex Taq HS (Takara, Shiga, Japan), 2 µL of 10×Ex Taq Buffer, 1.6 µL of dNTP Mixture, 1 µL of forward and reverse primer (10 µM), 12.3 µL sterilised distilled $H_2O$ and 2.0 µL DNA template. The PCR thermal conditions were as follows: 2 min at 94°C and 30 cycles of [30 s at 94°C, 25 s at 58°C and 60 s at 72°C] and 5 min at 72°C. Then PCR amplicons were purified using Sera-Mag SpeedBeads Carboxylate-Modified Magnetic Particles (Hydrophobic) (Cytiva, MA, USA).

(b) The mtDNA sequence including partial Cyt *b* (1,191 bp) of *N. sieboldii* was amplified using the following primer pairs: L14724 (5′-TGA CTT GAA RAA CCA YCG YYG-3′; Palumbi et al., 1991) and H15915 (5′-ACC TCC GAT CTY CGG ATT ACA AGA C-3′; Aoyama et al., 2000). The PCR was performed in a 15-µL total volume of reaction mixture containing 3.5 µL of GoTaq Green Master Mix (Promega, Tokyo Japan), 1.5 µL of forward and reverse primer (5 µM), 3.5 µL sterilised distilled $H_2O$ and 1.0 µL DNA template. The PCR thermal conditions were as follows: 2 min at 95°C and 30 cycles of [30 s at 95°C, 30 s at 48°C and 60 s at 72°C] and 5 min at 72°C. Then PCR amplicons were purified using Illustra ExoStar (GE Healthcare Japan, Tokyo, Japan).

For both species, the Sanger sequencing was performed using an automated DNA sequencer (GA3130xl; Applied Biosystems, Foster City, CA), amplification primers for each species and the BigDye Terminator Cycle sequencing FS Ready Reaction kit ver. 3.1 (Applied Biosystems). The obtained sequences were deposited in the DNA database DDBJ/EMBL/GenBank (accession No.: LC719969–LC719978 for *O. obscura*; LC718524–LC718549 for *N. sieboldii*).

### References


Aoyama, J., Watanabe, S., Ishikawa, S., Nishida, M., & Tsukamoto, K. (2000). Are morphological characters distinctive enough to discriminate between two species of freshwater eels,anguilla celebesensis andA. interioris? Ichthyological Research, 47(2), 157–161. https://doi.org/10.1007/BF02684236

Palumbi, S., Martin, A., Romano, S., McMillian, W., Stice, L., & Grabowski, G. (1991). The simple fool’s guide to PCR. Univ Hawaii, Honolulu.

Tsuji, S., Inui, R., Nakao, R., Miyazono, S., Saito, M., Kono, T., & Akamatsu, Y. (2022). Quantitative environmental DNA metabarcoding reflects quantitative capture data of fish community obtained by electrical shocker (p. 2022.04.27.489619). bioRxiv. https://doi.org/10.1101/2022.04.27.489619

Tsuji, S., Murakami, H., & Masuda, R. (2022). Analysis of the persistence and particle ssize distributional shift of sperm-derived environmental DNA to monitor Jack Mackerel spawning activity. Environmental Science & Technology. https://doi.org/10.1021/acs.est.2c01904
